# Mutations in TIC100 impair and repair chloroplast protein import and impact retrograde signalling

**DOI:** 10.1101/2022.01.18.476798

**Authors:** Naresh Loudya, Douglas P. F. Maffei, Jocelyn Bédard, Sabri Mohd. Ali, Paul Devlin, R. Paul Jarvis, Enrique López-Juez

## Abstract

Chloroplast biogenesis requires synthesis of proteins in the nucleocytoplasm and the chloroplast itself. Nucleus-encoded chloroplast proteins are imported via multiprotein translocons in the organelle’s envelope membranes. Controversy exists around whether a 1 MDa complex comprising TIC20, TIC100 and other proteins constitutes the inner membrane TIC translocon. The Arabidopsis *cue8* virescent mutant is broadly defective in plastid development. We identify *CUE8* as *TIC100*. The *tic100^cue8^* mutant accumulates reduced levels of 1 MDa complex components and exhibits reduced import of two nucleus-encoded chloroplast proteins of different import profiles. A search for suppressors of *tic100^cue8^* identified a second mutation within the same gene, *tic100^soh1^*, which rescues the visible, 1 MDa complex-subunit abundance, and chloroplast protein import phenotypes. *tic100^soh1^* retains but rapidly exits virescence, and rescues the synthetic lethality of *tic100^cue8^* when retrograde signalling is impaired by the *gun1* mutation. Alongside the strong virescence, changes in RNA editing and the presence of unimported precursor proteins show that a strong signalling response is triggered when TIC100 function is altered. Our results are consistent with a role for TIC100, and by extension the 1 MDa complex, in the chloroplast import of photosynthetic and non-photosynthetic proteins, a process which initiates retrograde signalling.

**One sentence summary:** Complementary mutations in TIC100 of the chloroplast inner envelope membrane cause reductions or corrective improvements in chloroplast protein import, and highlight a signalling role.

## Introduction

Chloroplast-containing photosynthetic eukaryotes sustain the biosphere. Chloroplast biogenesis is a complex process which in plants requires the involvement of 2000-3000 nucleus-encoded proteins and approximately 80 proteins encoded by the chloroplast’s own genome (Jarvis and López-Juez, 2013). The majority of proteins (those which are nucleus-encoded) need to be imported into the chloroplasts through the double-membrane envelope. This is achieved by the operation of protein import translocons at the outer and inner envelope membranes of chloroplasts – TOC and TIC, respectively (Jarvis and López-Juez, 2013; Nakai, 2018; Richardson and Schnell, 2020).

At the outer membrane TOC complex, subunits with GTPase activity act as receptors for the N-terminal targeting signals of chloroplast-destined polypeptides, and another subunit, TOC75, acts as a transmembrane import channel (Jarvis and López-Juez, 2013). At least two versions of the TOC complex exist, with different client specificities: one contains receptors with a preference for abundant photosynthetic pre-proteins, while the other favours import of house-keeping pre-proteins, like those involved in the chloroplast genetic machinery (Ivanova et al., 2004; Kubis et al., 2004; Jarvis and López-Juez, 2013). Targeted replacement of TOC receptor proteins has been revealed as a fundamental determinant of the development of photosynthetic or non-photosynthetic plastids and of plastid type transitions (Ling et al., 2012; Ling et al., 2019).

The identity of the inner membrane TIC components, in contrast, has been the subject of considerable debate. Initial studies identified an abundant 110 kDa protein (Kessler and Blobel, 1996; Lubeck et al., 1996), later named TIC110, and postulated to be a scaffold coordinating internal chaperones (Inaba et al., 2003) or, alternatively, the inner membrane import channel (Heins et al., 2002). Another candidate for the role of inner membrane channel is TIC20 (Chen et al., 2002; Kovacs-Bogdan et al., 2011), and a 1 MDa complex comprising nucleus-encoded TIC20 and at least three other proteins – including TIC100, TIC56 and chloroplast-encoded TIC214, but not including TIC110 – has been identified as a core channel-forming TIC complex (Kikuchi et al., 2013). An alternative form of TIC20 was shown to occur in root tissue, in the absence of the other components of the complex. However, the 1 MDa TIC complex model has proven controversial (de Vries et al., 2015; Nakai, 2015b; Bolter and Soll, 2017; Sjuts et al., 2017; Richardson and Schnell, 2020). Objections center around the low abundance of TIC20 compared to TIC110 (Vojta et al., 2004; Kovacs-Bogdan et al., 2011), the fact that the three additional proteins of the 1 MDa complex are absent in the grass family (Kikuchi et al., 2013; de Vries et al., 2015), the observation that the full-length version of TIC56 is dispensable in Arabidopsis (Kohler et al., 2015; Kohler et al., 2016; Schafer et al., 2019), and data pointing to other functions for TIC56 and TIC214 (Kohler et al., 2016; Schafer et al., 2019). The observation of a combined TOC-TIC “supercomplex” which includes TIC20 but also a small fraction of the total TIC110 (Chen and Li, 2017), leaves the issue of the nature of the channel unresolved.

Chloroplast development in flowering plants occurs exclusively in the light (Arsovski et al., 2012), because photoreceptors activate the expression of many genes for chloroplast-destined proteins (Cackett et al., 2021). A genetic screen for mutants in which light failed to activate the promoter of the *LHCB1*2* (*CAB3*) gene led to the identification of *CAB-underexpressed* (*cue*) mutants (Li et al., 1995; López-Juez et al., 1998). Among them, *cue8* exhibited a severe phenotype characterised by reduced plastid development in both dark and light conditions, and strongly impaired induction of (specifically) photosynthesis-associated genes by phytochrome photoreceptors (López-Juez et al., 1998; Vinti et al., 2005) linked to chloroplast-to-nucleus communication (Loudya et al., 2020). Seedlings of *cue8* display largely normal photomorphogenesis but have leaf rosettes with a virescent (slow-greening) phenotype. This virescence is due to a cellular correction phenomenon: an “anterograde” (nucleus-to-chloroplast) response which maintains a juvenile state of plastids, a delay in the transition from the pre-photosynthetic proplastid to differentiated chloroplast state, manifested in multiple physical and genetic features, and which allows for an eventual overcoming of the plastid defect (Loudya et al., 2020). This change is a response to “retrograde signals” (mediating chloroplast-to-nucleus communication), and can therefore be described as a “retro-anterograde correction”. Interestingly, evidence has recently accumulated pointing to an involvement of defects affecting the import of cytosol-synthesised proteins into chloroplasts (Wu et al., 2019; Tadini et al., 2020), or protein folding or quality control inside the organelle (Tadini et al., 2016; Wu et al., 2019; Tadini et al., 2020), in the initiation of changes in the cytosol. Such processes are broadly described as protein homeostasis or “proteostasis”, and their involvement triggers what can be described as a folding stress response, which may, in turn, cause retrograde signalling to the nucleus (Wu et al., 2019; Tadini et al., 2020). Indeed, GUN1, a chloroplast pentatricopeptide repeat protein which plays an important role in retrograde signalling, was shown to interact with chaperones involved in, or acting after, protein import, with its absence impairing import under specific conditions (Wu et al., 2019) and causing some depletion of components of the import machinery itself (Tadini et al., 2020).

We sought the molecular identity of the *CUE8* gene by positional cloning. We here report that *cue8* carries a missense mutation affecting TIC100, one of the components of the 1 MDa TIC complex. Furthermore, a genetic screen for suppressors of this mutant identified a second, intragenic mutation. Comprehensive analyses of both *tic100^cue8^* and the suppressed mutant (carrying two mutations in the same gene) demonstrated a significant role for this protein in chloroplast protein import, which is in turn consistent with such a role for the 1 MDa complex. In spite of the suppressed mutant’s recovery in import capacity, it retained a pronounced early virescence and exhibited strong genetic interaction with the loss of GUN1. These results, alongside others showing changes in RNA editing and gene expression and the likely occurrence of unimported polypeptides, highlighted the dramatic impact that changes in TIC100 have on chloroplast-to-nucleus communication.

## Results

### Mutation of *CUE8* leads to defects in plastid development in leaves and roots

The *cue8* mutant was previously identified following mutagenesis of the pOCA108 reporter-containing line (Figure 1A) (Li et al., 1995). We recently demonstrated that the virescent, slow-greening phenotype of *cue8* is associated with reduced chloroplast development in early cotyledons or very young leaf tissues (in which chloroplasts fail to fill the available cellular space), and by a gradual recovery of normal chloroplasts (Loudya et al., 2020). We wished to investigate whether the mutation impacts plastid development beyond leaves, widely across tissues, and so incorporated a plastid-targeted DsRed fluorescent protein (Haswell and Meyerowitz, 2006) into the wild type, and then introgressed the transgene into the *cue8* mutant. The fluorescence signal was substantially reduced in *cue8*, relative to wild type, both in cotyledon mesophyll cells and in roots, in which partially-developed chloroplasts were prominent in cells surrounding the central vasculature (Figure 1B and C). Accordingly, leaf development and root elongation were both reduced in the mutant (Supplemental Figure 1A and B). Supplementation of the growth medium with sucrose rescued the *cue8* root phenotype, in a dose-dependent manner but to an incomplete extent (Supplemental Figure 1). Thus, we concluded that CUE8 plays a role in plastid development in non-photosynthetic tissues, as well as in photosynthetic tissues.

**Figure 1.**
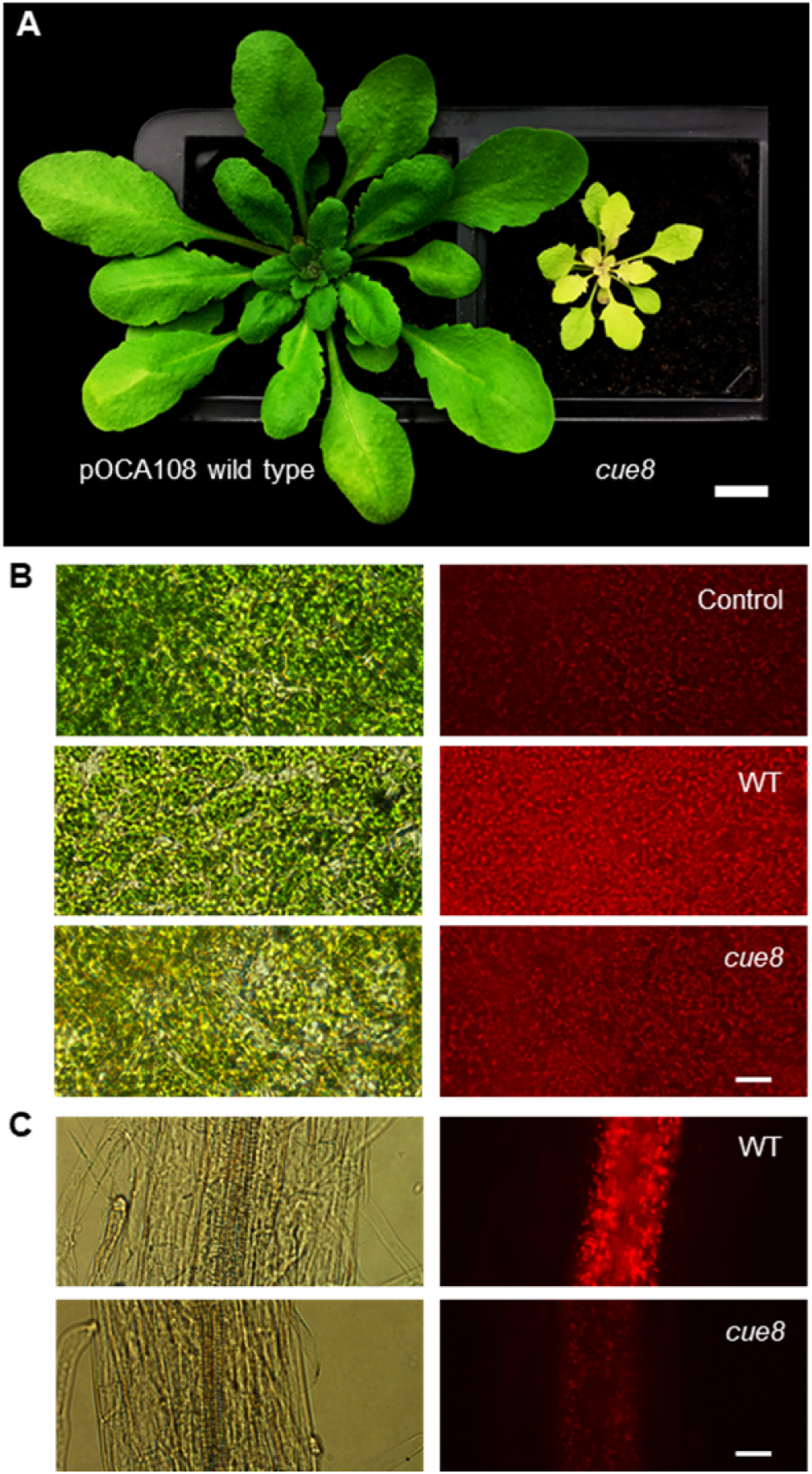
Mutation of *CUE8* causes a delay in plastid development in both aerial and root tissues. (**A**) Phenotype of 28-day-old pOCA108 wild type and *cue8* mutant plants. Scale bar 1 cm. (**B**) Mature cotyledon samples of plants without plastid-targeted dsRed (Control), or dsRed-containing pOCA108 wild type (5 days) or dsRed-containing *cue8* (6 days). (**C**) Root samples of equivalent seedlings. The same transgene was present in both genotypes, and images of WT and mutant were taken using the same exposure. Scale bar (**B** and **C**) 25 µm.

### Identification of the *CUE8* locus by linkage mapping

*cue8* and its wild-type progenitor (Li et al., 1995) are lines in the Bensheim ecotype of Arabidopsis (Figure 1A). We generated two mapping populations for *cue8* by performing outcrosses to both Landsberg-*erecta* (La-*er*) and Columbia (Col-0), to take advantage of ecotype polymorphisms (Supplemental Table 1). The *CUE8* locus was mapped to an 82 kb region of chromosome 5, containing 19 complete open reading frames (Figure 2A, see Materials and methods). A Transformation-competent Artificial Chromosome (TAC) covering 11 of those genes was able, when transformed into *cue8*, to complement the mutation (Supplemental Figure 2 and Supplemental Table 2). A combination of sequencing of individual candidate genes and assessment of the phenotypes of T-DNA knockouts (Supplemental Table 2) ruled out 10 of those genes, while we were unable to identify a viable homozygous mutant for AT5G22640 (only heterozygous T-DNA-containing plants were recovered). Sequencing of genomic DNA of *cue8* confirmed the presence of a mutation in this gene resulting in a G→R amino acid substitution at position 366, just outside one of the protein’s predicted Membrane Occupation and Recognition Nexus (MORN) domains (Takeshima et al., 2000) (Figure 2A, 5C, Supplemental Table 3). Transformation of the mutant with a wild-type (pOCA108) cDNA encoded by AT5G22640 under the control of a constitutive promoter also resulted in complementation (Figure 2B and C). Thus, we concluded that *CUE8* is AT5G22640, a gene identified previously as *EMB1211*, due to its embryo-lethal knockout mutant phenotype (Liang et al., 2010), and, most interestingly, as *TIC100* (Kikuchi et al., 2013), encoding a component of the putative 1 MDa TIC complex. We hereafter refer to the mutant allele, and the plant carrying it, as *tic100^cue8^*. Bearing in mind the nature of the *tic100^cue8^* amino acid substitution, as well as the mutant’s virescent phenotype, which contrasts with the loss of viability caused by a T-DNA insertion at this locus, we concluded that the *tic100^cue8^* is a hypomorphic allele, carrying a missense mutation which causes a partial loss-of-function of the gene.

**Figure 2.**
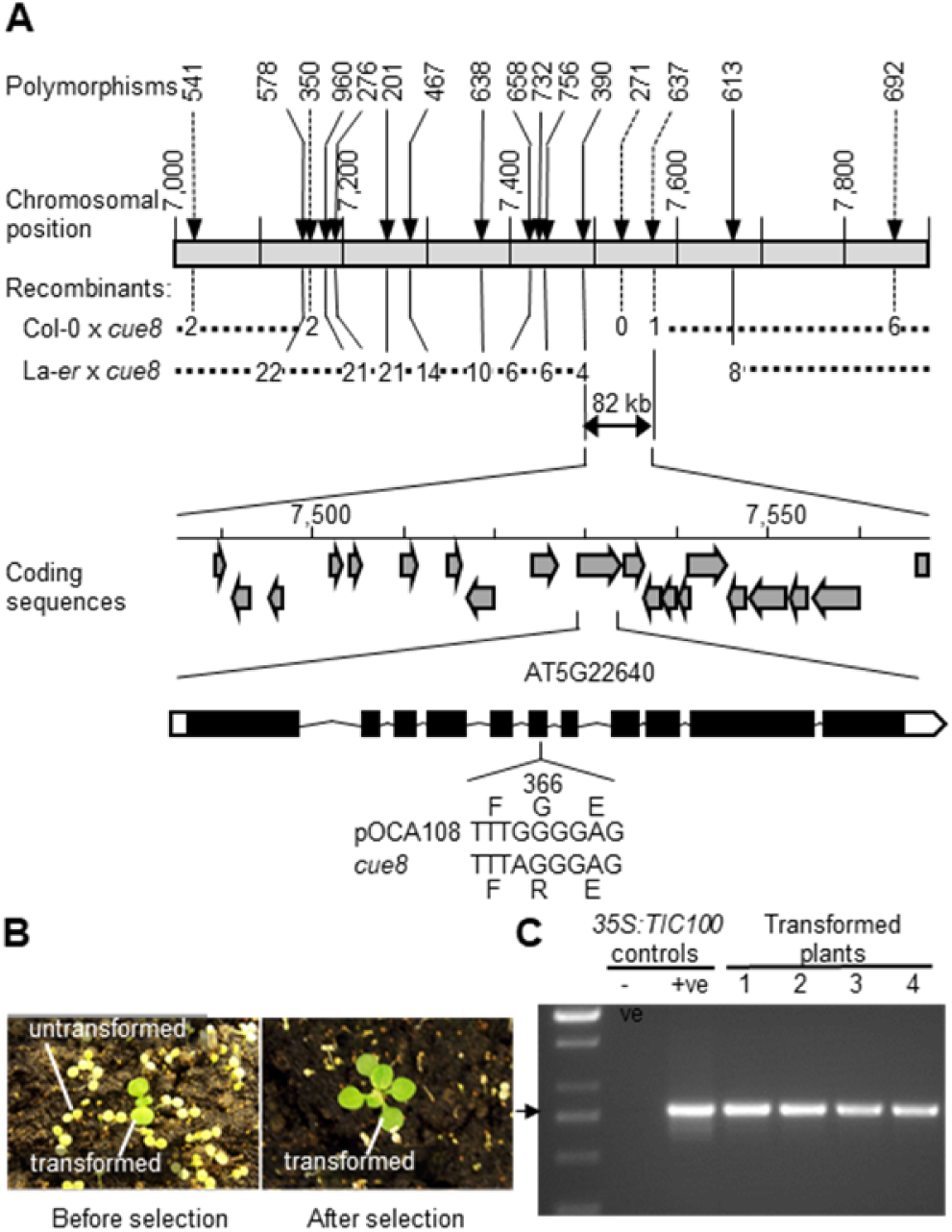
Cloning of *CUE8*. **(A)** Map-based cloning of the *CUE8* gene, AT5G22640, encoding *TIC100* / *EMB1211*. Upper panel: abbreviated name of the polymorphisms used for mapping, position along chromosome 5 (in kb), and number of recombinants at those positions identified in the indicated mapping populations. This identifies an 82 kb region, containing 19 open reading frames (middle panel). A combination of strategies (Supplemental Table 2) identifies AT5G22640 (*TIC100*) as the *CUE8* locus, whose exon/intron structure is shown. A point mutation (lower panel) results in a single amino acid substitution (G366R) in the TIC100 protein sequence. **(B)** Complementation of *cue8* with *35S:TIC100*, carrying a *TIC100* cDNA under the control of a 35S promoter. Plants shown before and after the selection of transformants. **(C)** Diagnostic PCR confirming the presence of the *35S:TIC100* transgene in complemented plants. Positive (+ve) control, plasmid DNA harbouring the construct. Negative (-ve) control, DNA from plant prior to transformation.

Several chloroplast protein import components have previously been shown to have a preferential role in the import of either abundant, photosynthetic proteins or less-abundant, but essential, plastid housekeeping proteins (Jarvis, 2008). Having identified *CUE8*, we compared, using publicly-available data (Schmid et al., 2005), its developmental expression with that of genes representative of those two functions. The *CUE8*/*TIC100* gene exhibited (Supplemental Figure 3) a combined expression pattern: high like *LHCB2.1* in photosynthetic tissues, while also high like *TOC34* in those tissues rich in meristematic cells, such as the root tip. Results of a search for co-regulated genes, using two different algorithms (Supplemental Datasets 1 and 2) were also consistent with *CUE8*/*TIC100* being involved early (for example, together with transcription and translation, pigment synthesis and protein import functions) in the biogenesis of photosynthetic as well as non-photosynthetic plastids.

### Reduced protein import rate and partial loss of the 1 MDa complex in *tic100^cue8^* chloroplasts

Taking advantage of the opportunity afforded by the partial loss of function of TIC100 in *tic100^cue8^*, we carried out *in vitro* import assays with chloroplasts isolated from well-developed seedlings of the mutant, using a photosynthetic protein precursor, the Rubisco small subunit (SSU). Four independent experiments, using developmentally-comparable wild-type and mutant plants (Figure 3A), revealed that *cue8* mutant chloroplasts import less than one-third of the amount of pre-protein than the equivalent number of wild-type chloroplasts (Figure 3B, C).

**Figure 3.**
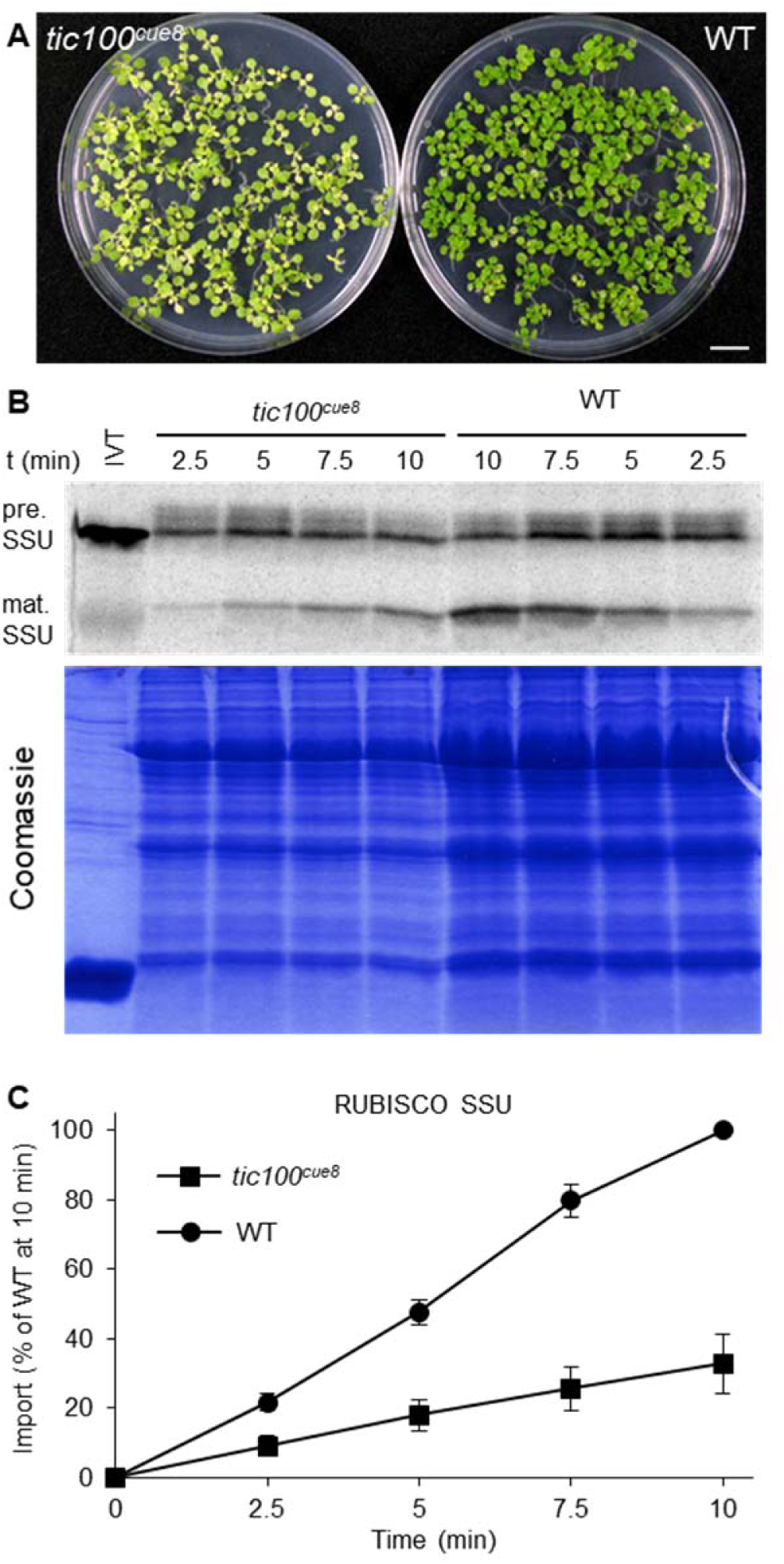
Chloroplasts of *tic100^cue8^* exhibit reduced protein import rates. **(A)** 13-day-old wild-type and developmentally-comparable 17-day-old *tic100^cue8^* mutant seedlings, used to isolate chloroplasts for import assays. Seedlings were grown on 0.5% sucrose. Scale bar: 1 cm. **(B)** Phosphor screen image of import reactions of *in vitro*-translated RUBISCO SSU polypeptide (IVT), carried out with equal total numbers of chloroplasts isolated from the seedlings above. Samples were taken 2.5, 5, 7.5 or 10 min after the start of the reaction. The import reaction converts the precursor (pre.) into mature polypeptide (mat.) of reduced size. Results from one representative experiment. A Coomassie-stained total protein gel corresponding to the same experiment is also shown. **(C)** Quantification of the amount of mature protein at each time point, normalised relative to the amount of mature protein in WT after 10 min of import. Average values from four independent experiments. Error bars represent s.e.m. Values for *tic100^cue8^* were significantly different to those of WT at every time point (Student’s t-test, 2-tailed, p<0.05).

To understand more clearly the basis for the protein import deficiency in the mutant, we analysed the levels of several translocon components by immunoblotting. Equal amounts of total chloroplast proteins were loaded per lane. Band intensities for some translocon components, in both the outer (TOC75) and inner (TIC110, TIC40) envelope membranes, were elevated by about a third in the *tic100^cue8^* lanes (Supplemental Figure 4). This reflected the fact that *tic100^cue8^* chloroplasts are, to varying extents, less developed internally and contain reduced amounts of the major photosynthetic proteins, including Rubisco and LHCB (López-Juez et al., 1998), relative to wild type, leading to the relative overloading of envelope components in the *tic100^cue8^* samples when using equal protein amounts (this also explains the slightly lower amounts of total protein in the *tic100^cue8^* samples following normalisation according to equal chloroplast numbers in Figure 3B). Crucially, in spite of this, the level of TIC100 polypeptide was reduced to between one quarter and one eighth that in the wild type (Supplemental Figure 4) on an equal total chloroplast protein basis, or less than one eighth when normalized to another envelope protein, TIC40. The decrease in TIC100 abundance in the mutant was linked to reductions in the levels of the other components of the 1 MDa complex (TIC20 and the additional TIC56 and TIC214; Supplemental Figure 4), to between 25 and 50% of wild-type levels when expressed relative to TIC40. These observations are consistent with the notion that these proteins associate, with the very substantial loss of *tic100^cue8^* preventing others from accumulating normally.

### Identification of a suppressor mutation of *tic100^cue8^*

A search for suppressor mutations of the *ppi1* mutant, defective in the TOC33 subunit of the outer envelope translocon, led to the identification of SP1, a ubiquitin ligase which remodels the import complexes to control protein import and plastid development (Ling et al., 2012; Ling et al., 2019). We sought to deepen our understanding of inner envelope translocation processes by searching for suppressors of the *tic100^cue8^* mutation. Screening of mutagenised M2 populations for increased levels of greening led to the identification of a mutant with a dramatic phenotype, which we named *suppressor of tic100 1*, *soh1* (Figure 4A). Backcrossing of *tic100^cue8^ soh1* into the *tic100^cue8^* parent resulted in 100% of the F1 (62 seedlings) showing a phenotype which was intermediate between that of the parents but closer to the *soh1* phenotype (Supplemental Figure 5A); while self-pollination of the F1 plants yielded 75% (554 out of 699, Chi-squared p=0.19) seedlings with suppressed phenotype, among which about a third displayed a marginally larger seedling phenotype (these plants represented a quarter of the total F2 population: 158 out of 699, Chi-squared *p*=0.21). These data indicated that *soh1* is a gain-of-function mutation that improves greening and growth, and which has a semi-dominant character.

**Figure 4.**
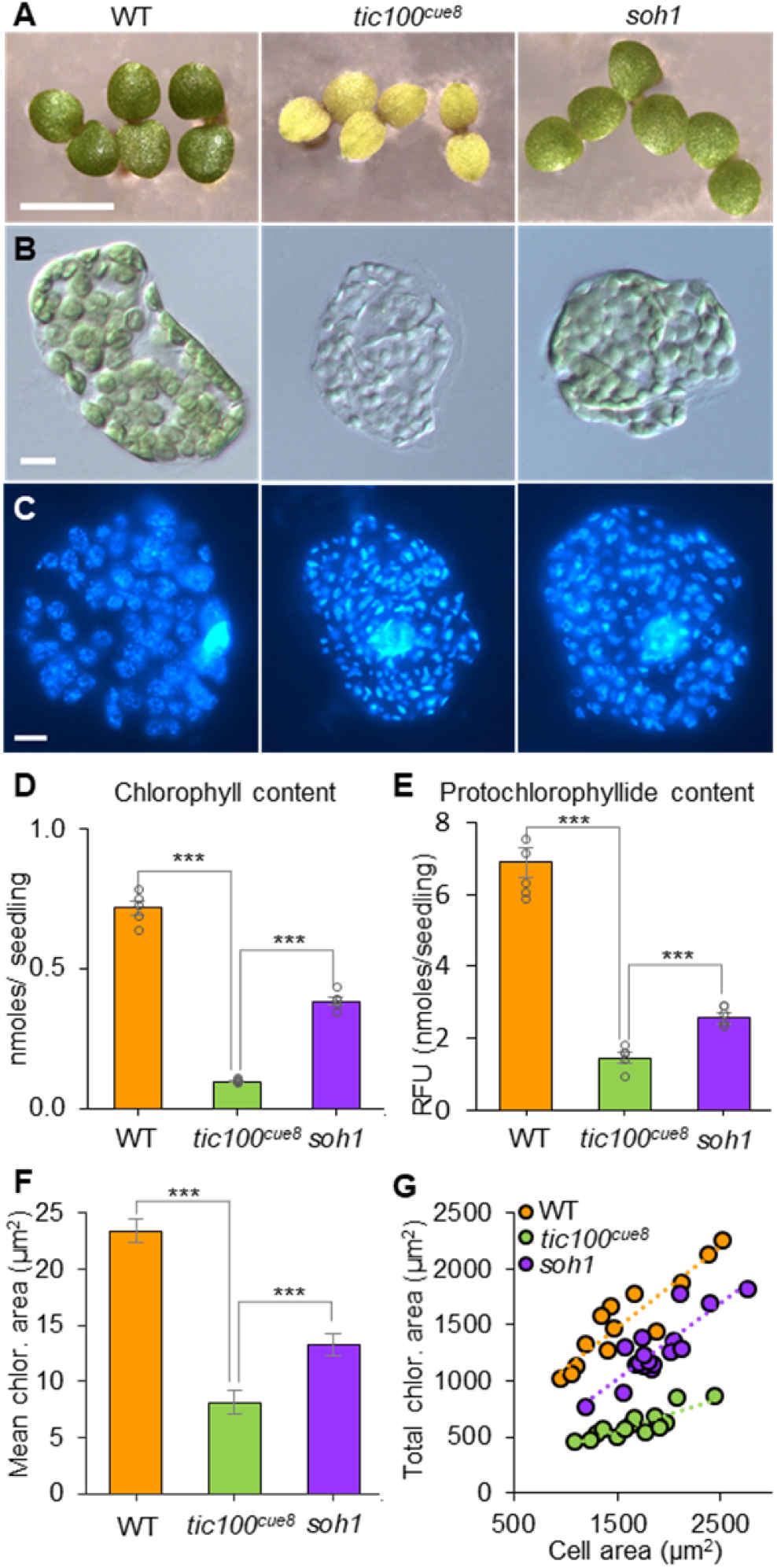
Identification of *soh1,* a suppressor mutant of *tic100^cue8^*, and phenotype of young cotyledon cells and their chloroplasts in wild type, *tic100^cue8^* and *soh1* seedlings. (A) Five-day (WT) and six-day (*tic100^cue8^* and *soh1*) seedlings. Scale bar: 5 mm. (B) Individual cells of the three genotypes, of seedlings equivalent to those in A, observed under DIC microscopy, displaying the different degrees of cell occupancy by chloroplasts. (C) Individual cells observed fluorescence microscopy following DAPI-staining of double-stranded DNA, revealing both the nuclei and the presence and density of nucleoids in individual chloroplasts. Scale bar (B and C) 10 µm. (D) Chlorophyll content per seedling for seedlings identical to those in A. (E) Protochlorophyllide content per seedling (RFU, relative fluorescence units) of 5-day-old seedlings of the three genotypes. (F) Mean area of individual chloroplasts in cells equivalent to those in B. (G) Total plan area of chloroplasts in a cell plotted against cell plan area, for the three genotypes, including regression lines of best fit. The presented values are means, and the error bars (in D, E) show s.e.m. from five biological replicates, each with at least 5 seedlings, or (in F) at least 10 individual chloroplasts from each of at least 13 individual cells total, obtained from at least four different cotyledons per genotype. For all panels, asterisks above lines denote comparisons under the lines: ***P < 0.001 (2-tailed Student’s t-test).

We have recently shown that the virescent phenotype of *tic100^cue8^* is caused by early chloroplasts in very young cotyledons being small and unable to fill the available cellular space (Loudya et al., 2020). In this regard, chloroplasts of 6-day-old *soh1* seedlings were much closer to those in the wild type (Figure 4B). Consequently, plastid DNA nucleoids, tightly packed in *tic100^cue8^* as previously reported (Loudya et al., 2020), appeared much less dense in *soh1* (Figure 4C). Moreover, total chlorophyll of light-grown seedlings, protochlorophyllide of dark-grown seedlings, the average size of individual chloroplasts, and the mesophyll cellular occupancy by chloroplasts (the chloroplast index), which were all reduced dramatically in *tic100^cue8^*, were largely restored in *soh1* seedlings (Figure 4D-G).

### Identification of the gene carrying the *soh1* mutation

We generated a mapping population by backcrossing the suppressor mutant as originally identified (*tic100^cue8^ soh1*, Bensheim ecotype) to the unmutagenised *tic100^cue8^* parent (also in the Bensheim ecotype). F2 seedlings of unsuppressed, *tic100^cue8^* phenotype were used for mapping by SHORT READS sequencing (SHOREmap) (Schneeberger et al., 2009), as described in Supplemental Figure 5B-C. Mapping of the *soh1* mutation identified a region of chromosome 5 (Supplemental Figure 5D) spanning seven genes with mutations in the open reading frame, and one of these was *TIC100*. Sanger sequencing confirmed the presence in the *soh1* mutant of both the original *tic100^cue8^* mutation and a second mutation, which we provisionally named *tic100^soh1^* (Figure 5A-C). Constitutive expression under the 35S promoter of a *tic100^cue8 soh1^* cDNA carrying both of these mutations in the *tic100^cue8^* mutant plants resulted in T1 plants showing a suppressed phenotype (Figure 5E); the genotyping of these plants confirmed the presence of both *tic100^cue8^* (from the endogenous gene) and *tic100^cue8 soh1^* alleles (from the transgene). In contrast, constitutive expression of the *tic100^cue8^* cDNA under the same promoter in *tic100^cue8^* plants produced only *tic100^cue8^* phenotypes (Supplemental Figure 6), demonstrating that it was the second mutation, and not overexpression of the gene, that caused the suppression effect. Therefore, we concluded that the second mutation was indeed responsible for the suppressed phenotype, and we hereafter refer to the *tic100^cue8 soh1^* double mutant as *tic100^soh1^* (Figure 5A).

**Figure 5.**
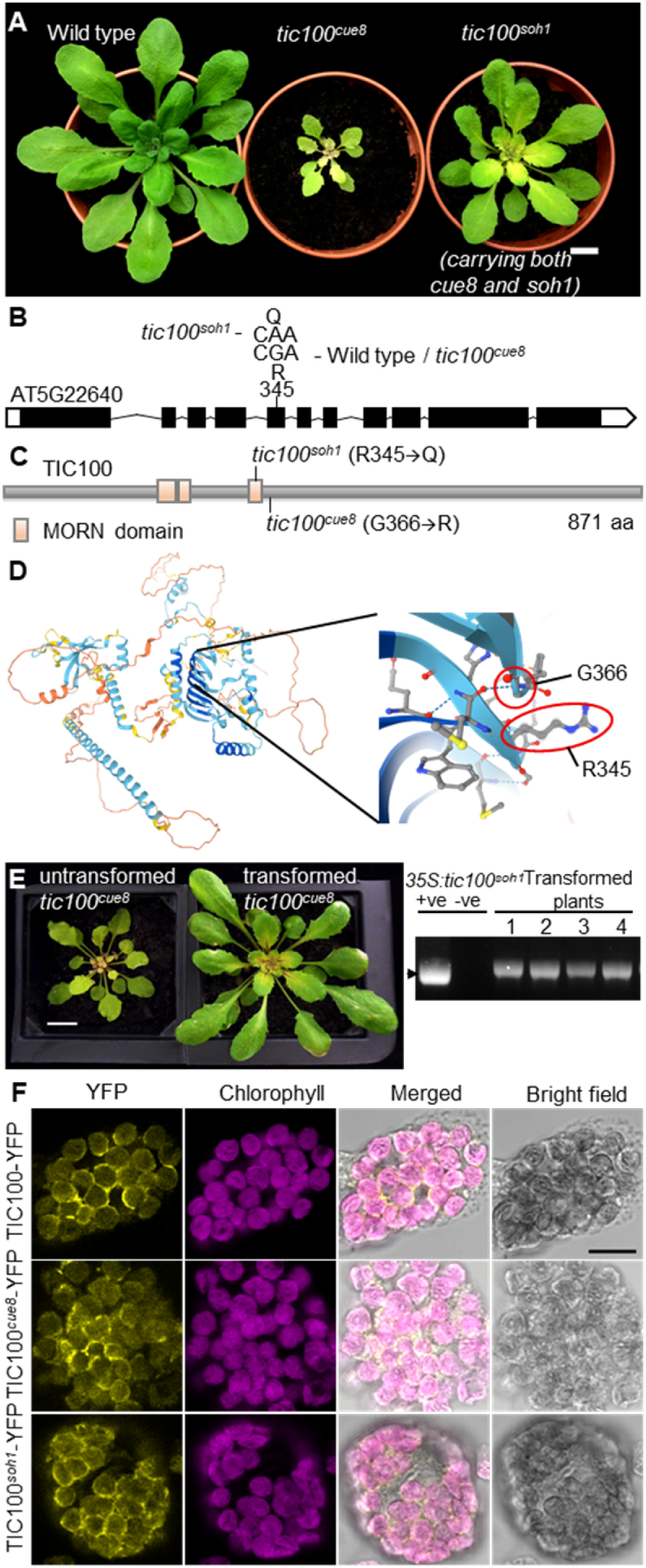
Cloning of the intragenic *suppressor of TIC100* (*soh1*) mutation, phenocopying and localization of TIC100. **(A)** Soil grown plants of wild type pOCA108, *tic100^cue8^* (*tic100^cue8^*) and suppressed *tic100^cue8^ soh1* double mutants (*tic100^soh1^*) are shown at 28 days of age. Scale bar: 1 cm. **(B)** Second missense mutation (and resulting substituted aminoacid position) in the genomic sequence of the *TIC100* gene present in the *tic100^soh1^* mutant and absent in *tic100cue8* or its wild type parent. **(C)** Model of the domain structure of the TIC100 protein, indicating the position of the single mutation present in *tic100^cue8^*, or the two mutations present in the *tic100^soh1^* double mutant. The MORN domains occupy positions 219-239, 243-257 and 337-352. **(D)** TIC100 protein structure prediction by Alphafold, showing the position of the wild type aminoacids affected by the *tic100 ^cue8^* and *tic100 ^soh1^* mutations. Light and dark blue represent regions of confident and highly-confident prediction respectively. **(E)** Phenocopying of the suppressor *soh1* mutant by transformation of the single *tic100 ^cue8^* mutant with an over-expressed, double-mutated *tic100 ^soh1^* coding sequence driven by the 35S promoter (as seen in 11 independent T1 plants, 4 shown). Plants shown at 30 days of age. Scale bar: 1cm. Gel on the right confirms the genotype of the transformed plants. “+ve”: positive genotyping control (bacterial plasmid). **(F)** Localisation of the TIC100 protein, in its wild type, TIC100^cue8^ and TIC100^soh1^ (double-mutated) forms, to the chloroplast periphery in transfected protoplasts. Wild-type protoplasts were transfected with constructs encoding wild-type and mutant forms of TIC100, each one tagged with a C-terminal YFP tag. The protoplasts were analysed by confocal microscopy. Images represent results of at least two independent experiments (at least 40 protoplasts per genotype) showing the same result. Scale bar: 10 μm.

Analysis of the predicted domain structure of the TIC100 protein by searching in the Interpro domains database showed that the initial *tic100^cue8^* mutation occurred immediately outside the C-terminus of the third of three MORN domains, introducing a basic arginine residue in place of a neutral glycine. Conversely, the *tic100^soh1^* mutation replaced an arginine residue, within the third MORN domain (20 amino acids upstream of the *tic100^cue8^* substitution), with a neutral glutamine residue (Figure 5B, C). Three-dimensional protein structure prediction by the recent, breakthrough AlphaFold algorithm (Jumper et al., 2021) indeed showed the aminoacids affected by the two substitutions to lie in very close proximity in space, in regions of confidently predicted structure (Figure 5D), at one end of a large, highly confidently predicted β-sheet region which includes the MORN domains.

### TIC100^cue8^ and TIC100^soh1^ proteins retain localisation at the chloroplast periphery

Previous biochemical analyses identified the TIC100 protein as part of the 1 MDa complex in the inner envelope membrane with a proposed role in pre-protein import (Kikuchi et al., 2013; Chen and Li, 2017; Richardson et al., 2018). In view of the protein import defect of the mutant, described above, we asked whether the *tic100^cue8^* mutation interferes with the localisation of the TIC100 protein, and whether such an effect might in turn be alleviated by the *tic100^soh1^* mutation. Therefore, we constructed YFP fusion versions of the TIC100 protein in its wild-type and two mutant forms (the second carrying both mutations). Transient overexpression of the fusions in protoplasts resulted in some accumulation of all three proteins in the cytosol of the cells (Supplemental Figure 7), which interfered with assessment of chloroplast envelope association. However, in protoplasts in which rupture of the plasma membrane eliminated the background cytosolic protein (which we interpret to be mislocalised owing to overexpression), TIC100 was clearly observed at the periphery of chloroplasts, possibly with a small amount of intra-organellar signal; this is consistent with the previous biochemically-determined localisation. Significantly, neither of the two mutations altered this character (Figure 5F), and so we concluded that the mutations affect a property of TIC100 other than its localisation.

### *tic100^soh1^* corrects the protein import defect caused by *tic100^cue8^*

Next, we asked whether the basis for the suppression of the *tic100^cue8^* phenotype in the *tic100^soh1^* mutant was a correction of the plastid protein import defect described earlier. In these experiments, the import of the photosynthetic SSU pre-protein into equal numbers of chloroplasts isolated from wild-type, *tic100^cue8^* single-mutant, and *tic100^soh1^* double-mutant plants (Figure 6A, B) was measured. On this occasion, the results demonstrated a reduction of protein import in *tic100^cue8^* to approximately 55% of the wild-type level; the smaller reduction in import seen here, relative to Figure 3, was attributed to the slightly greater extent of development of the mutant plants on a higher sucrose concentration in the medium (Supplemental Figure 1). Notably, protein import into the suppressed mutant chloroplasts was restored almost completely to wild-type levels (over 90%) (Figure 6C, E).

**Figure 6.**
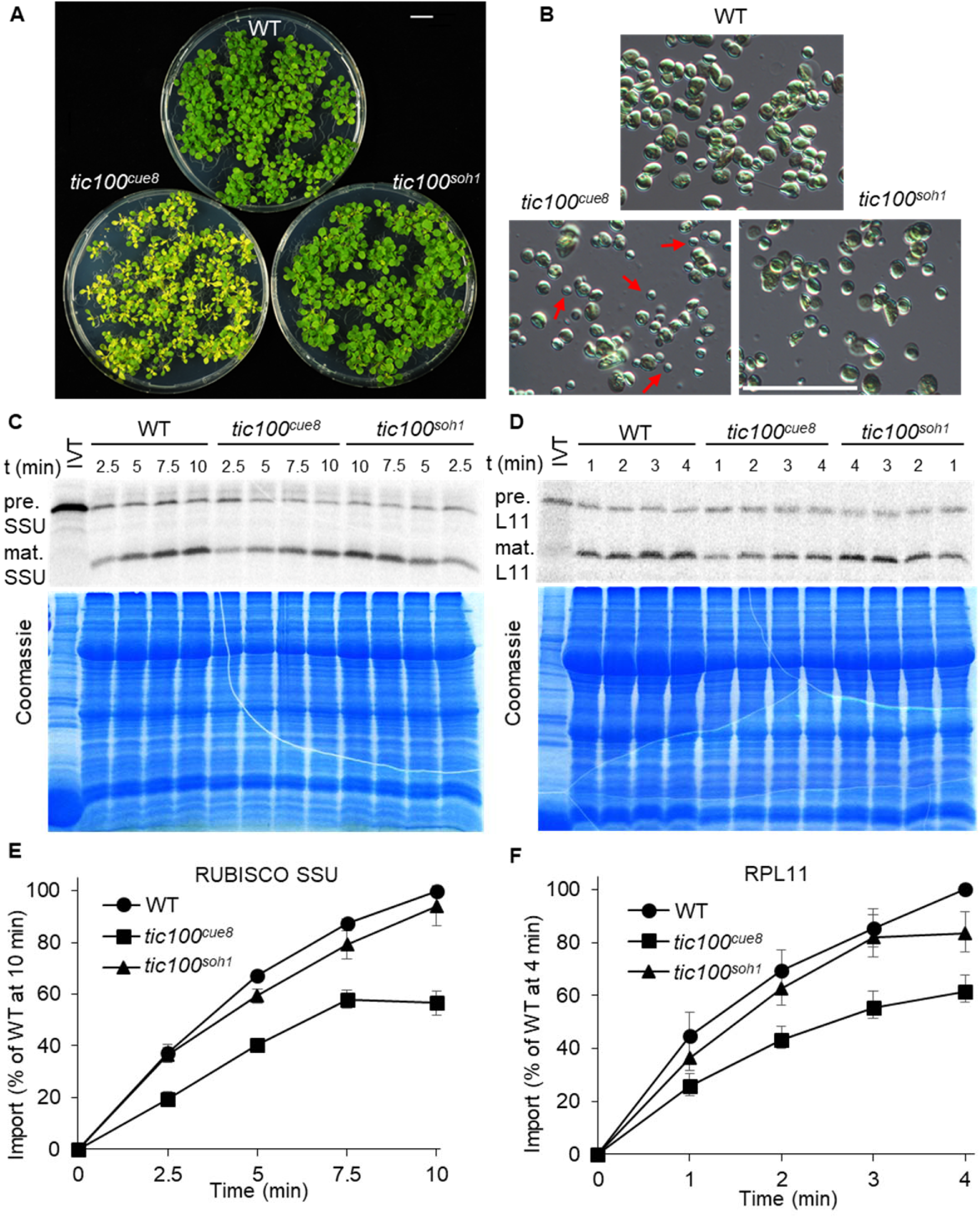
Chloroplasts of *tic100^cue8^* exhibit reduced import of a photosynthetic and a housekeeping pre-protein, with different age-dependent import profiles, and both defects are suppressed by the second mutation in *tic100^soh1^.* (A) 13-day-old wild-type, and 17-day-old *tic100^cue8^* and *tic100^soh1^* mutant, seedlings used to isolate chloroplasts for the import assays. Seedlings were grown on 1% sucrose. Scale bar: 1 cm. (B) Examples of chloroplast populations used for the *in vitro* protein import assays. Occasional small chloroplasts in *tic100^cue8^* are indicated with red arrows. Scale bar: 50 µm. (C) Phosphor screen image of import reactions of *in vitro*-translated Rubisco small subunit (SSU) polypeptide (IVT), carried out with equal numbers of *tic100^cue8^* and wild-type chloroplasts. Samples were taken at the indicated times after the start of the reaction. The import reaction converts the precursor (pre.) into the mature (mat.) polypeptide of reduced size. Results from one representative experiment are shown. The lower panel shows the corresponding Coomassie-stained total protein gel of the same experiment. (D) Import of RPL11 into equal numbers of wild type, *tic100^cue8^*and *tic100^soh1^* chloroplasts. Upper and lower panels as in C. Values at every time point were significantly different for *tic100^cue8^* relative to WT, and for *tic100^soh1^* relative to *tic100^cue8^*. (E) Quantitation of at least four independent protein import assays as that shown in C, from four separate chloroplast populations obtained from at least four groups of independently grown plants. The presented values are means, and the error bars show s.e.m. (F) Quantitation of at least four independent import assays, as that shown in D. Values at all time points for SSU and 2, 3 and 4 minutes for RPL11 were significantly different for *tic100^cue8^* relative to WT, and for *tic100^soh1^* relative to *tic100^cue8^* (*p*<0.05, 2-tailed Student’s t-test).

To further assess the role of TIC100 in relation to functionally different proteins, we additionally tested the import of the housekeeping plastid RPL11 protein (50S plastid ribosomal subunit protein). This also allowed us to assess whether reductions in import capacity in *tic100^cue8^* chloroplasts could simply be an indirect consequence of differences in the stage of development of mutant chloroplasts, since SSU and RPL11 have been shown to be preferentially imported by chloroplasts of younger or older leaves, respectively (Teng et al., 2012). To the contrary, we obtained very similar results for RPL11 to those we had obtained for SSU (Figure 6D, F).

Overall, these protein import data (which were observed across four independent experiments per pre-protein) revealed a clear import defect in *tic100^cue8^* for two proteins which display different developmental stage-associated import profiles (Teng et al., 2012), and which use different types of TOC complexes (Jarvis and López-Juez, 2013; Demarsy et al., 2014). This was consistent with the gene expression profile of *TIC100* (Supplemental Figure 3). Notably, this was also consistent with the fact that SSU preprotein was previously shown to physically associate during import with several components of the 1 MDa complex, including TIC100, while RPL11 preprotein was found to associate with TIC214 (Kikuchi et al., 2018), Moreover, our results revealed a very pronounced correction of the import defects seen in *tic100^cue8^* chloroplasts in the *tic100^soh1^* suppressed mutant.

### *tic100^soh1^* restores levels of 1 MDa protein components in *tic100^cue8^*

Given the strong reductions in levels of 1 MDa TIC complex components (but not of other inner or outer envelope proteins) seen in *tic100^cue8^* mutant chloroplasts (Supplemental Figure 4), we asked whether the *tic100^soh1^* mutation had corrected the accumulation of components of the 1 MDa complex. Immunoblot analyses indicated that this was indeed the case (Figure 7A-C). Analysis of TIC100 protein using the same chloroplast preparations as used for import assays in Figure 6 showed partial restoration of the level of this protein in the double mutant. Furthermore, qualitatively similar trends to those seen for TIC100 were observed for TIC56 and TIC214, but we were unable to quantify TIC20 here due to very limited availability of the corresponding antibody. In contrast, no such protein level reduction in *tic100^cue8^*, or restoration in *tic100^soh1^*, was observed for control housekeeping, non-membrane proteins (HSP70, RPL2); in fact envelope proteins unrelated to the 1 MDa complex (TIC110, TIC40 and TOC75) appeared elevated in *tic100^cue8^*, consistent with an enrichment of envelope proteins per unit total chloroplast protein, as discussed earlier, and accordingly returned to normal apparent levels in *tic100^soh1^*.

**Figure 7.**
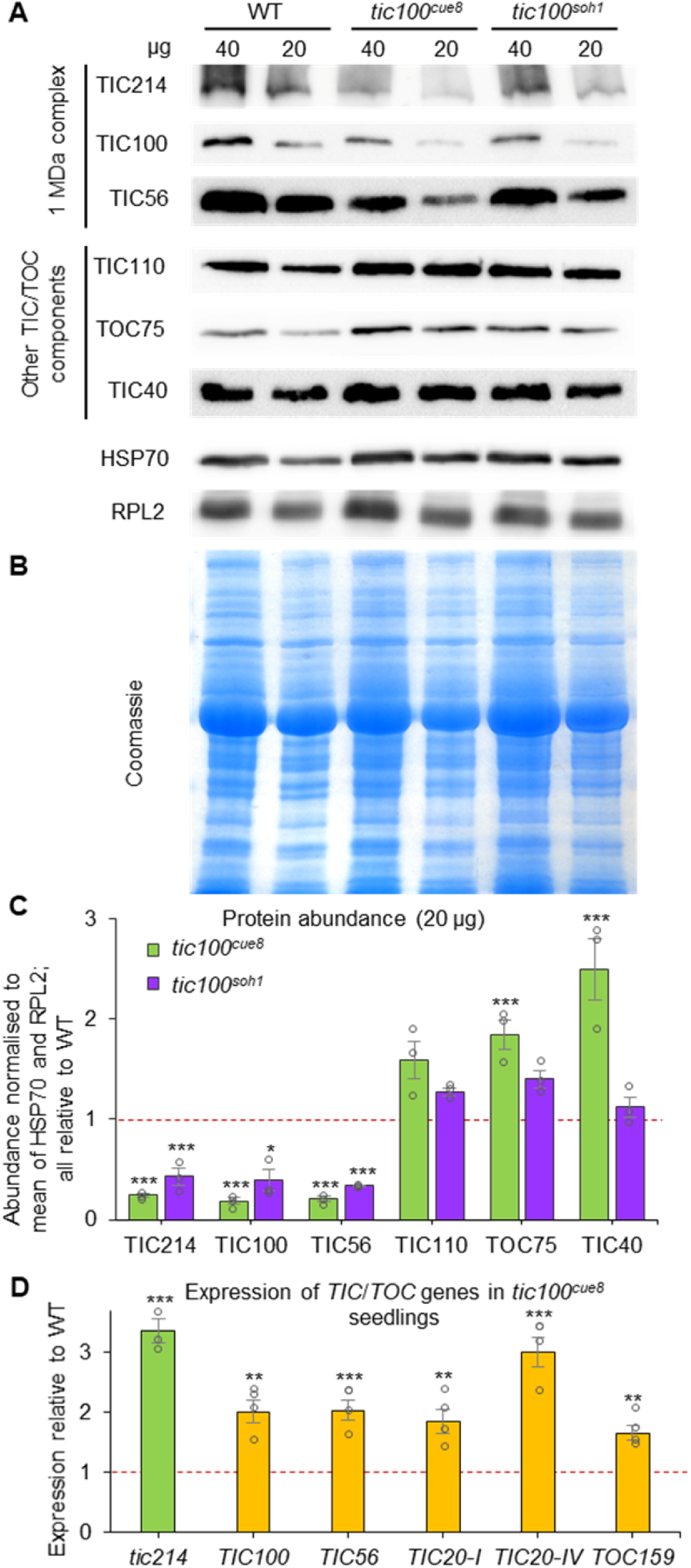
Chloroplasts of *tic100^cue8^* display decreased levels of 1 MDa complex proteins specifically, and this defect is suppressed by the second mutation in *tic100^soh1^*. (A) Immunoblot analysis of total chloroplast proteins from preparations from wild type (13-day-old), and *tic100^cue8^* and *tic100^soh1^* (17-day-old), seedlings (see Figure 6). The amount of proteins (µg) loaded is indicated above each lane. The antibodies used for the detection of components of the 1 MDa complex (TIC56, TIC100 and TIC214) or other chloroplast envelope proteins (TOC75, TIC40 and TIC110) are indicated. Note the reduced amounts of components of the 1 MDa complex, which is apparent despite the increased loading of envelope proteins (as revealed by the levels of other polypeptides) specifically in the *tic100^cue8^* samples. Very limited antibody availability precluded probing the chloroplast protein extracts for the levels of TIC20. (B) Coomassie-stained total protein gel of the same experiment. (C) Quantitation of protein abundance from an analysis of the 20 µg samples in three independent experiments, relative to the mean of HSP70 and RPL2 in each sample, all expressed relative to wild-type protein levels. The presented values are means, and the error bars show s.e.m. Asterisks represent significance of difference of each mutant relative to WT (ANOVA followed by Dunnett’s test). (D) Expression, measured by quantitative real-time RT-PCR, of *TIC*/*TOC* genes in *tic100^cue8^* seedlings similar to those analysed in Figure 3, measured relative to expression in wild-type seedlings. Note *tic214* is chloroplast-encoded. The presented values are means, and the error bars show s.e.m. of three RNA samples (biological replicates), each with two technical replicates. Asterisks represent significance of difference between mutant and WT: *P < 0.05, **P < 0.01, ***P < 0.001 (2-tailed Student’s t-test). Dotted lines represent protein levels (C) or expression (D) in WT.

It could be argued that the reduced accumulation of the 1 MDa TIC might have been an indirect result from a retrograde signalling impact of the *tic100^cue8^* mutation, leading to reduced nuclear gene expression and synthesis of translocon components. Such an explanation is highly unlikely given that we have previously observed elevated, not reduced, expression of nuclear and chloroplast-encoded genes for early-expressed, plastid housekeeping proteins in the mutant (Loudya et al., 2020). This is part of its “pre-photosynthetic, juvenile plastid” phenotype. In fact, we confirmed here that the expression of nucleus-encoded genes for 1 MDa TIC components and, especially, of the plastid-encoded *tic214* gene, were all elevated in *tic100^cue8^* (Figure 7D); the expression of the control *TOC159* gene was also elevated. Interestingly, and as previously observed for other elements of the juvenile plastid phenotype (Loudya et al., 2020), a substantial component of the retro-anterograde correction did not involve GUN1 action, since it occurred in *tic100^cue8^* even in the absence of GUN1 (Supplemental Figure 8).

Moreover, the expression of *TIC20-IV*, which encodes an alternative form of TIC20 that functions independently of the 1 MDa complex (Kikuchi 2013), was also elevated in *tic100^cue8^* (Figure 7D). Thus, we concluded that accumulation of subunits of the TIC 1 MDa complex is reduced by the *tic100^cue8^* mutation at the posttranscriptional level, that this occurs in spite of the attempted “retro-anterograde correction” at the gene expression level brought about by retrograde signalling, and that the accumulation of TIC 1 MDa complex subunits is partially restored by the *tic100^soh1^* mutation. Furthermore, a potential compensatory effect of the *tic100^cue8^* mutation, increasing the expression of *TIC20-IV* encoding an alternative TIC20 form that acts independently of the 1 MDa complex, is apparent.

### Interplay between tic100 mutations and retrograde signalling

We previously observed that while the single *tic100^cue8^* mutation resulted in virescence, the simultaneous loss of GUN1 (which in itself causes partial uncoupling of nuclear gene expression from the state of the plastid) was incompatible with survival – i.e., combination of the *tic100^cue8^* and *gun1* mutations resulted in synthetic seedling lethality in the double mutants (Loudya et al., 2020). To further investigate the extent of suppression in *tic100^soh1^*, we analysed *tic100^soh1^ gun1* triple mutants. In contrast to *tic100^cue8^ gun1* albino, eventually lethal mutants, the *tic100^soh1^ gun1* triple mutants were very pale but not lethal (Figure 8A), and indeed they could survive and produce seeds entirely photoautotrophycally under low light conditions. In other words, the synthetic lethality of *tic100^cue8^ gun1* double mutants was suppressed by *tic100^soh1^*.

**Figure 8.**
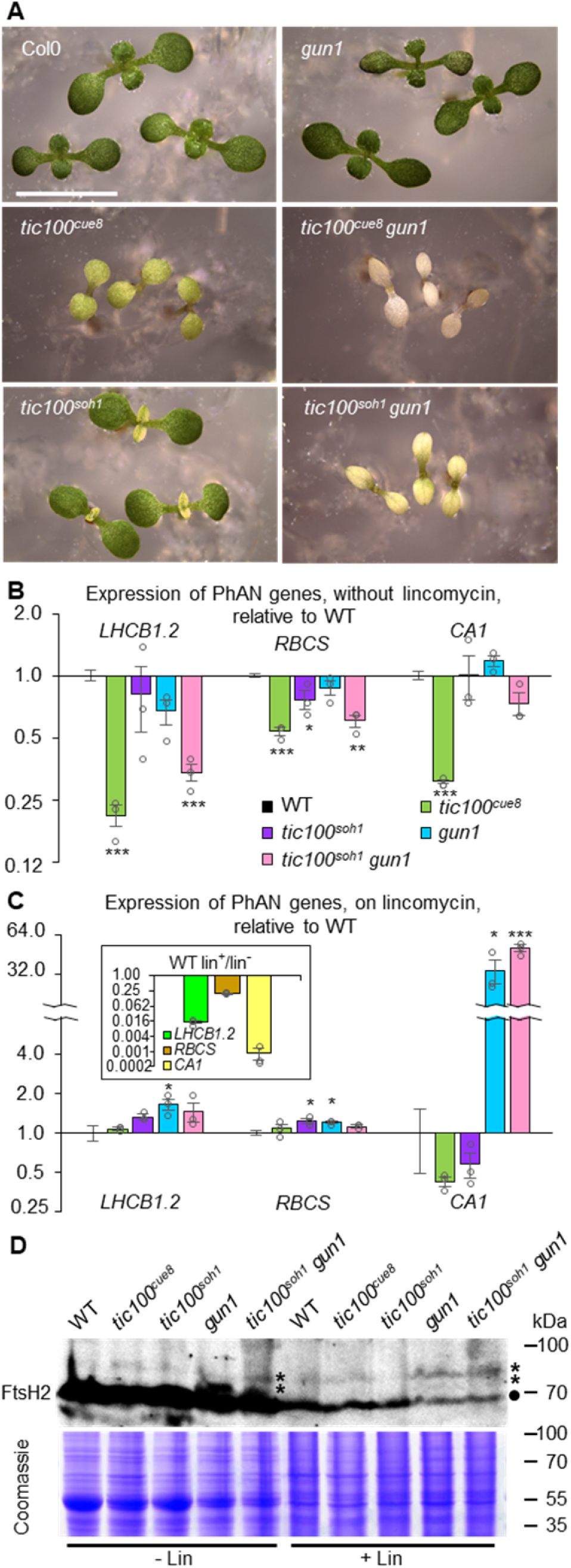
The *tic100^soh1^* mutation suppresses the synthetic seedling lethality of *tic100^cue8^* in combination with *gun1*, while *gun1* retains its “uncoupled” phenotype and unimported FtsH2 precursors can be detected in the *tic* mutants. (A) Phenotype of 7-day-old seedlings of the genotypes indicated. *tic100^cue8^ gun1* double mutants exhibit eventual seedling lethality. Scale bar: 5 mm. (B, C) Photosynthesis-associated nuclear (PhAN) gene expression in the absence (B) or presence (C) of lincomycin in the genotypes indicated. Asterisks denote significance of difference between mutant and WT as indicated for Figure 7D. The inset compares the phenotype of the wild type grown on lincomycin with that in its absence. Differences for all three genes exhibited P < 0.001. (D) Immunoblot detection of higher molecular weight bands, previously shown to represent unimported, cytosolic precursors (asterisks) of chloroplastic FtsH2 (circle), in the presence or absence of lincomycin, in the genotypes indicated.

Evidence has recently accumulated for a role of altered organelle proteostasis in chloroplast retrograde signaling (Tadini et al., 2016; Wu et al., 2019; Tadini et al., 2020). The clear virescence exhibited by *tic100^soh1^*, and its strong, albeit reduced, genetic interaction with loss of GUN1, led us to ask whether retrograde signalling was in any way altered by impairment or recovery of TIC100 function. Two scenarios were in principle possible: In the first, a strong response of reduced photosynthesis-associated nuclear gene (PhANG) expression might be observed when TIC100 function is reduced, resulting in the virescence of *tic100^cue8^* and even *tic100^soh1^* mutants, and the very low PhANG expression in *tic100^cue8^* (Vinti et al., 2005),. In this scenario, GUN1 function, mediating retrograde signalling, would remain fundamentally unchanged in the mutants. In the second scenario, the impairment of protein import occurring in *tic100^cue8^*, but barely so in *tic100^soh1^*, would result in the accumulation of unimported proteins in the cytosol of the mutants, and this in turn, as proposed by Wu and coworkers (Wu et al., 2019), would itself cause elevated PhANG expression in spite of the chloroplast damage, i.e., a *genomes uncoupled* (*gun*) phenotype. We examined these two possible, contrasting scenarios by quantifying transcript levels of *LHCB1.2*, *RBCS* and *CA1* – the first two genes classically monitored in retrograde analyses, and the third gene showing one of the greatest extents of reduction by treatment with lincomycin (Koussevitzky et al., 2007) – in the presence of a chloroplast translation inhibitor which triggers a dramatic loss of PhANG expression. We exposed to lincomycin seedlings of each *tic100* mutant, and of the *tic100^soh1^ gun1* triple mutant. We did not examine *tic100^cue8^ gun1*, as such albino seedlings become impossible to select in the presence of the antibiotic from the segregating population in which they occur. The results of this analysis (Figure 8B-C) are clearly consistent with the first scenario: reductions in PhANG expression were strong in the absence of lincomycin in *tic100^cue8^ gun1* double and in the *tic100^soh1^ gun1* triple mutants (Figure 8B), and mild in *tic100^soh1^*. The reductions in the double mutant were not due to the *gun1* mutation having lost its associated “uncoupled” phenotype, but rather to the very strong chloroplast defect (manifested as a greening defect) in the double. This was shown by the fact that in *tic100^soh1^ gun1* PhANG expression, particularly that of *CA1*, was clearly uncoupled, i.e., much less reduced by lincomycin than it was in the wild type (Figure 8C). We concluded that, as anticipated, *tic100^cue8^* is not a *gun* mutant. We also concluded that the capacity of GUN1 to initiate retrograde communication remains strong in *tic100^soh1^*, and that it can therefore explain the retained, pronounced virescence.

PhANG expression changes, in particular in seedlings grown on lincomycin, have been attributed to the presence of unimported precursor proteins, which can be detected in whole-cell and cytosolic extracts of such antibiotic-treated plants (Wu et al., 2019; Tadini et al., 2020). Such precursors have also been detected in *gun1* even in the absence of antibiotic (Tadini et al., 2020), and it was reasonable to speculate that they might be also detectable in the *tic100* mutants. We examined this through immunoblot analysis of the FtsH2 protein, a thylakoid-associated chaperone for which high molecular weight (HMW) bands have been previously observed (Tadini et al., 2020) using the same antibody. The HMW bands were previously shown to represent cytosolic, unimported precursors by cellular fractionation and by the construction and examination of FtsH2-GFP fusion proteins. Our results (Figure 8D) confirmed that the level of mature FtsH2 protein is much reduced in extracts of lincomycin-treated seedlings. Furthermore, the antibody could detect the presence of the same HMW bands (unimported protein) in extracts of seedlings grown in the presence of the antibiotic, and also in those of mutant seedlings in its absence, an observation which is consistent with a chloroplast protein import defect in both cases.

### Editing of chloroplast mRNA is altered in *tic100^soh1^ gun1* seedlings in a manner consistent with the juvenile plastid phenotype

RNA editing occurs for many chloroplast transcripts, and it has been previously demonstrated that growth of seedlings on norflurazon (which blocks carotenoid synthesis) or lincomycin, and the concomitant disruption of chloroplast development, results in changes in the extent of editing of different chloroplast transcripts (Kakizaki et al., 2012). Importantly, such changes, involving both increases and decreases in editing, were also observed in plants defective in the TOC159 outer membrane translocon receptor (Kakizaki et al., 2012). They were also seen in *tic100^cue8^* and, even more dramatically, *tic100^cue8^ gun1* (Loudya et al., 2020); and they involved increased editing of *rpoC1*, encoding a subunit of the chloroplast RNA polymerase, and decreased of *ndhB*, encoding a photosynthetic electron transport protein. We interpreted such changes, not as evidence of a direct role of CUE8 (TIC100) in editing, but as part of the “juvenile plastid” phenotype of the mutants. That interpretation is consistent with the existence of two phases of organelle biogenesis: an early, “plastid development” phase (photosynthesis-enabling but pre-photosynthetic, involving expression of the chloroplast genetic machinery), and a later, “chloroplast development” phase involving photosynthetic gene expression (as seen particularly clearly in developing cereal leaves: (Chotewutmontri and Barkan, 2016; Loudya et al., 2021). We asked whether this aspect of the *tic100^cue8^* phenotype had also been suppressed by *tic100^soh1^*. This was indeed the case (Figure 9): the extent of editing of *rpoC1* and *ndhB* transcripts was indistinguishable between *tic100^soh1^* and the wild type (or *gun1*). In contrast, editing was increased for *rpoC1*, and reduced for *ndhB*, in the *tic100^soh1^ gun1* double mutant, which we again interpret as a more juvenile state of chloroplast development caused by the combination of the mild loss of import capacity and the simultaneous loss of GUN1 function.

**Figure 9.**
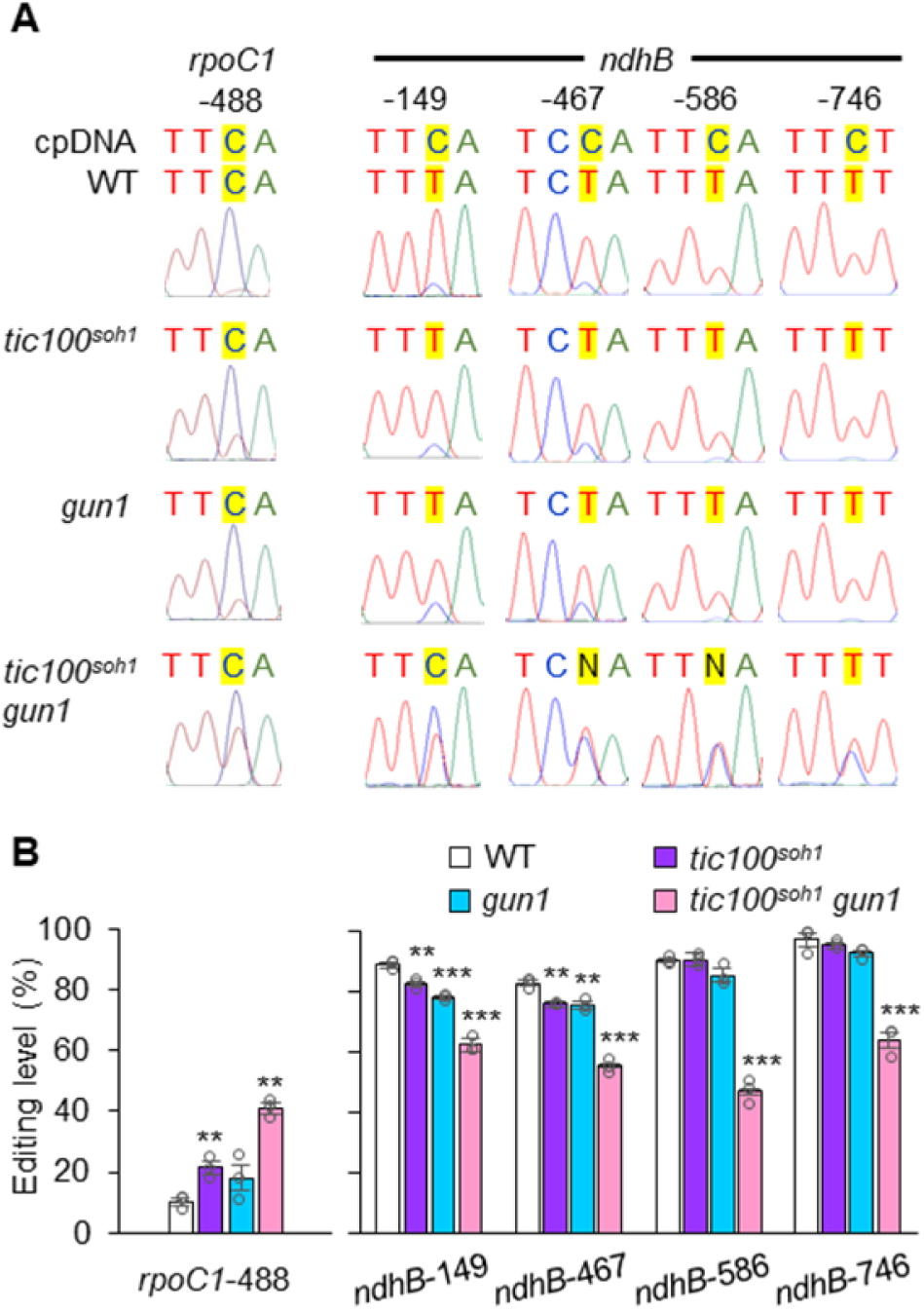
Editing of chloroplast mRNA is increased for the transcripts of *rpoC1* and decreased for those of *ndhB* in *tic100^soh1^ gun1* seedlings. (A) Representative sequence electropherograms of cDNA generated from cDNA of the genotypes indicated. The original, genomic cpDNA sequence is indicated at the top. (B) Quantitation of the degree of editing of the two plastid mRNAs shown in (A), averaged for three independent cDNA preparations from different seedling samples per genotype. Asterisks denote significance of difference between mutant and WT as indicated for Figure 7D.

## Discussion

The nature of the proteins imported into plastids clearly determines plastid type and functions, and the identity of the import receptors even influences whole-plant performance in the face of stress. This underscores the importance of questions concerning the exact composition of the protein translocons. Indeed, the identity of the inner membrane TIC machinery has been the subject of much debate, particularly concerning the identity of the channel and the import motor (de Vries et al., 2015; Nakai, 2015b; Bolter and Soll, 2017; Sjuts et al., 2017; Schafer et al., 2019; Nakai, 2020; Richardson and Schnell, 2020). The serendipitous identification of *cue8* as a hypomorphic allele of *TIC100* provided an opportunity to begin addressing one such area of controversy. Knockout mutants of *TIC100*/*EMB1211* (a component of TIC 1 MDa complex) suffer from severe embryo development defects, leading to very early seedling lethality (Liang et al., 2010; Kikuchi et al., 2013). Attempts have previously been made to address the role of the 1 MDa complex by assessing the import characteristics of the seedling-lethal knockout mutant *tic56*-1, by proteomically analysing chloroplast-targeted proteins in such seedlings (Kohler et al., 2015). However such analysis is indirect and cannot, by definition, reveal differences in import rate, since seedlings are examined after lengthy *in vitro* culture. A second mutant, *tic56*-3, expresses a truncated TIC56 protein and has a mild phenotype, allowing the execution of *in vitro* assays to determine chloroplast protein import rates (Kikuchi et al., 2013; Kohler et al., 2015). Perhaps owing to different growth or chloroplast preparation conditions, those experiments did (Kikuchi et al., 2013) or did not (Kohler et al., 2015) observe reduced import rates in the mutant. In fact, one analysis reported very low levels of 1 MDa complex subunits overall in the *tic56*-3 mutant, and failed to detect TIC100 at all (Schafer et al., 2019), while in the same mutant TIC100 had previously been readily observable (Kikuchi et al., 2013). Understandably, it has been difficult to reach consensus based on data obtained from such genotypes.

An alternative approach may help to provide a resolution. Informative *in vitro* protein import rate assays were readily performed in the current study using the chloroplasts of *tic100^cue8^*, which are severely impaired in TIC100 accumulation due to the *tic100^cue8^* mutation. Our data demonstrated a clear reduction in the efficiency of protein import into mutant chloroplasts, which cannot be explained simply by their reduced size or extent of development. Importantly, we have previously observed a physical and gene expression phenotype consistent with a juvenile state of plastid development in *tic100^cue8^* plastids (Loudya et al., 2020), and it is well known that plastids of young plants generally achieve greater, not inferior, import efficiencies (Dahlin and Cline, 1991). The use of two different preproteins with different age-dependency of import profiles (Teng et al., 2012; Chu et al., 2020), both of which exhibited reduced import in *tic100^cue8^*, indicated that the import reduction was not an indirect consequence of an altered developmental stage of the mutant organelles. On the contrary, the reduced protein import seen in *tic100^cue8^* chloroplasts provided a clear molecular explanation for the mutant’s strongly virescent phenotype; while the increased rate of protein import seen in *tic100^soh1^* chloroplasts explained the reduction of this virescence. We should stress that the suppressor mutation present in *tic100^soh1^* is unlikely to, in itself, have a positive impact on TIC100 function. It rather removes a native positive charge present in very close proximity to the additional positive charge introduced by the *tic100^cue8^* mutation. The suppressor mutation is therefore likely to have removed an electrostatic repulsion present in the *tic100^cue8^* mutant protein, which allowed the conformation of TIC100 to return to a mildly-impaired state, close to its native one.

The link between the abundance of TIC100 and that of its putative 1 MDa complex partners, in both *tic100^cue8^* and the intragenic suppressor mutant *tic100^soh1^*, is consistent with the existence of this complex. Furthermore, the protein accumulation profile data revealed a correlation between protein import rates and 1 MDa complex protein levels: decreased (*tic100^cue8^*) or increased (*tic100^soh1^*) protein import rates were observed as subunits of the complex decreased (*tic100^cue8^*) or increased (*tic100^soh1^*), in a genetically-determined manner. The data further showed that some loss of the subunits, as seen in *tic100^soh1^*, can be tolerated with minimal deleterious impact on protein import at the chloroplast-containing seedling stage; an equivalent phenomenon could potentially explain some of the previous conflicting observations on *tic56*-3. Thus, our data are consistent with the existence of the TIC100-containing 1 MDa complex and with it having a role in protein import. Our results also confirm the need for the complex for the normal import of different types of pre-proteins, exemplified by Rubisco SSU, a model photosynthetic protein, and RPL11, a non-photosynthetic protein. Nonetheless, we emphasise that the data presented here do not in any way rule out an important role for TIC110 in protein import; for example, this protein may function at a later stage in the import process, by providing a scaffold for the coordination of stromal chaperones as previously proposed (Inaba et al., 2003; Jarvis and López-Juez, 2013; Tsai et al., 2013; Richardson and Schnell, 2020).

Among the difficulties raised concerning the proposed role of the 1 MDa complex in protein import, one is of a phylogenetic nature: the absence of several of the complex’s main components from the grass family of monocots (de Vries et al., 2015; Bolter and Soll, 2016). In contrast, recent evidence strongly supports the function of the complex in a chlorophyte alga (Ramundo et al., 2020). It has been argued that an alternative form of the TIC20 translocon may operate in grasses, and that this utilises orthologues of TIC20-IV which is expressed and presumably active in roots of Arabidopsis (Kasmati et al., 2011; Nakai, 2015a). In this light, it is worth noting that the developmental impairment seen in roots of *tic100^cue8^* was not as pronounced as that observed in shoot meristem-derived tissues, suggesting a reduced need for TIC100 in the roots compared to shoots. Interestingly, we observed in *tic100^cue8^* a three-fold increase in expression of *TIC20-IV*. This may be a compensation for the loss of function of TIC20-I that occurs in the absence of its 1 MDa partners, as the level of expression of *TIC20-I* is four-fold higher than that of *TIC20-IV* in emerging seedling cotyledons and six-fold higher in young leaves (Klepikova et al., 2016). TIC20-IV, independent of the 1 MDa complex, might have also escaped the obvious posttranscriptional regulation we observed for components of the complex in the *tic100^cue8^* mutant. Nevertheless we do not believe that TIC20-IV could fully compensate for defects in 1 MDa complex, given the lethality of complex knockout mutants. It is apparent that there are some unquestionable differences between grasses and the majority of other flowering plant groups in terms of chloroplast biogenesis in leaves. For example, while the greening of dicot leaf primordia is noticeable from the very youngest stages, the greening of grass leaves occurs over a longer developmental period. Differences have also been seen for otherwise important components involved in organelle biogenesis; for example grass family genomes contain genes for one chloroplast-targeted and one mitochondrion-targeted RNA polymerase, but no gene for a protein targeted to both, whereas dicots do carry such a dual-targeted enzyme (Borner et al., 2015).

At present we can only speculate on the specific role of TIC100 within the 1 MDa complex. The nature of the *soh1* mutation highlighted the importance of at least one of the three recognised MORN domains in the protein. Such domains have been shown to be important in other proteins for membrane association and for association with specific lipids (Takeshima et al., 2000). Previous experiments (Kikuchi et al., 2013) have revealed that this protein, like TIC56, most likely occupies an intermembrane space position associated with the inner envelope membrane, while TIC20 (the protein with channel properties (Kovacs-Bogdan et al., 2011)) and TIC214 are integral membrane proteins. Our study also confirmed a localisation consistent with envelope association for TIC100, and one may speculate that this association is partly mediated by the MORN domains. However, our confocal microscopy analysis did not reveal any change in localisation in the TIC100^cue8^ and TIC100^soh1^ mutant proteins. Therefore, we interpret that the mutations either affect or rescue some other aspect of TIC100 function, such as interactions with other members of the 1 MDa complex to promote complex stability.

A role (possibly an additional one) for TIC56 in chloroplast ribosome assembly has been reported, and mild translation inhibition phenocopies many aspects of the *tic56*-3 phenotype (Kohler et al., 2016). These observations have been raised as an objection against the involvement of the TOC56 protein, and by extension of the 1 MDa complex, in protein import. However, intriguingly, such translation inhibition also phenocopies the phenotype caused by the loss the receptor protein TOC159, which has an extremely well established role in import (Kohler et al., 2016). Reduced accumulation of two import-related but chloroplast-encoded proteins, TIC214 and Ycf2/FtsHi – the latter a subunit of a putative import motor (Kikuchi et al., 2018) – might explain such similarities.

Intriguingly, *tic100^soh1^* exhibited almost complete rescue of protein import rates and of greening in fully-developed leaf tissue, and yet it still displayed a pronounced, early virescence phenotype. This virescence is consistent with a strong early impact on plastid-to-nucleus communication (a strong early retrograde signal, should the signal be a negative regulator of photosynthetic gene expression; or a strong absence of one, should the signal be a positive regulator). Indeed, the *tic100^soh1^ gun1* double mutant was very severely greening-deficient, although not seedling lethal. Combined loss of TOC159 and GUN1 has also been reported to have severe consequences (in this case, seedling lethality) (Kakizaki et al., 2012). A number of important connections between GUN1 and the chloroplast protein import apparatus have recently been uncovered. *gun1* mutants lose subunits of the import translocons, including TIC100, in response to mild inhibition of chloroplast translation, to a greater extent than the wild type does (Tadini et al., 2020). A 50% reduction in levels of import translocon subunits in *gun1*, observed on antibiotic-free medium (Tadini et al., 2020), could have made seedlings somewhat more sensitive to the defects brought about by the *tic100* mutations. However, that cannot fully explain the very strong genetic interactions between the respective mutations, reaching seedling lethality in the case of *tic100^cue8^ gun1*. *gun1* also causes mild but synergistic decreases in import rates in a mutant defective in chloroplast proteostasis, and GUN1 physically associates with chloroplast chaperones that act in protein import (Wu et al., 2019). Both of those studies (Wu et al., 2019; Tadini et al., 2020) demonstrated the accumulation of unimported preproteins in the wild type in the presence of lincomycin, and in *gun1* even in its absence. We observed HMW bands of FtsH2 that likely represent such unimported preproteins, and could particularly detect such bands in young seedlings of the mutant *tic100* genotypes, further supporting the import defects occurring in them. (Wu et al., 2019) also observed the emergence of a cellular folding stress response in the cytosol of chloroplast proteostasis mutants, consistent with the presence of such unimported proteins. We should note, though, that our evidence is consistent with the accumulation of unimported preproteins in the cytosol playing a major signalling role which results in the reduction, not the maintenance, of PhANG expression. The reduction in PhANG expression in response to impairment of TIC100 function particularly in *tic100^cue8^*, or to exposure to lincomycin, both of which may or do lead to preprotein accumulation, is consistent only with such a negative role. Therefore, our data do not support this aspect of the previously proposed model (Wu et al., 2019), of a role for increased accumulation of preproteins in elevating PhANG expression and therefore being the cause of the *gun* phenotype in the *gun1* mutant. We also did not observe a *gun* phenotype in *tic100^cue8^*, nor, in fact, was a *gun* phenotype observed in *toc33* (*ppi1*), *toc75-III*-3 and *tic40*-4 mutants (Wu et al., 2019). Our data support a loss of protein import at the inner envelope bringing about a reduction in PhANG expression and triggering a retro-anterograde delay in chloroplast development which requires GUN1 and, by allowing gradual correction of the defect, has adaptive value (Loudya et al., 2020). Our previous and current data on RNA editing in *tic100* mutants also support the notion that the shifts in degree of RNA editing occurring for different chloroplast transcripts also constitute part of such a retro-anterograde correction.

Taking these observations together, it is becoming apparent that the status of organelle protein import – particularly at the inner envelope membrane, mediated by the 1 MDa TIC complex – and protein homeostasis are critically interlinked with intracellular communication, and monitoring them is a critical function of chloroplast retrograde signalling, and of the GUN1 protein specifically. According to our observations, and consistently with previous ones (Kubis et al., 2003), impaired protein import reduces PhANG expression. How GUN1 relays information of changes in import status appears to remain unresolved, and warrants future exploration.

## Methods

### Plant material and growth conditions

The *Arabidopsis thaliana cue8* mutant (López-Juez et al., 1998; Vinti et al., 2005) and its wild type pOCA108, in the Bensheim ecotype, have been described. The *gun1*-1 mutant, in the Col-0 ecotype, was previously described (Susek et al., 1993).The *cue8* and *soh1* mutations were backcrossed into Col-0 for double mutant analysis as described (Loudya et al., 2020). The generation of the *soh1* mutant is described below. Plants were grown in soil under 16 h photoperiods and a fluence rate of 180 µmol m^-2^ s^-1^, and seedlings grown *in vitro* in MS media supplemented with 1% sucrose, unless otherwise indicated (Supplemental Figure 1) under continuous white light, at a fluence rate of 100 µmol m^-2^ s^-1^, as previously described (López-Juez et al., 1998; Loudya et al., 2020). Genotyping of the individual mutations (following gene identification), individually or for double mutant generation, used PCR followed by restriction digestion (Supplemental Table 4).

### Analysis of plastid development

Wild type and *cue8* lines carrying the DsRed reporter gene targeted to chloroplasts (Haswell and Meyerowitz, 2006) were identified following a cross and selected to homozygosity. Cotyledons and roots from *in vitro*-grown seedlings (7-day-old) were mounted on slides and observed using a Nikon (Kingston upon Thames, UK) Eclipse NI fluorescence microscope, x20 Plan Fluor objective and Texas Red filter block. Cotyledons of non-DsRed, negative control seedlings were examined to confirm that the majority of the fluorescence signal was attributable to the DsRed plastid reporter (Figure 1). Fluorescence images of the same type of tissue used identical exposure conditions.

Five-day-old wild type and six-day-old mutant seedlings were fixed (Figure 4) in 3.5% glutaraldehyde and subject to cell separation in 0.1M EDTA, pH9, prior to observation in a differential interference contrast Nikon Optiphot-2 microscope. Cells (n=13-18) of four independent cotyledons per genotype were observed, with cell plan areas, chloroplast number and individual area calculated as described (Loudya et al., 2020).

### Analysis of root development

Seedlings were cultured *in vitro*, under the conditions described in Supplemental Figure 1, images taken and root length quantified using ImageJ (ImageJ.net) software.

### Map-based cloning of *cue8*

Two mapping populations were generated following a *cue8* x La-*er* cross. In one, F2 mutant plants were selected (genotype *cue8*/*cue8*); in another, wild type (WT) plants were selected, grown to maturity and their progeny individually scored to identify plants without *cue8* progeny (genotype *+*/*+*). A third mutant mapping population was generated following a *cue8* x Col-0 cross. 344, 557 and 619 plants were selected respectively (total 1520 plants) in the three mapping populations. Plants were examined at polymorphic markers 541 and 692 (Col-0 population) or 576 and 613 (La-*er* populations). DNA was extracted from pools of 3-4 plants, was examined and, if a recombination event identified, individual plants were retested to identify the recombinant. Other polymorphisms between Col-0 and La-*er* (TAIR, www.arabidopsis.org) were developed as polymorphic markers by designing flanking primers for PCR amplification and differential enzyme digestion (Supplemental Table 1), and screening Col-0, La-*er* and pOCA108 genomic DNA. pOCA108 sequence was more frequently found to be polymorphic against La-*er* (13/20) than against Col-0 (7/20). Genes in the region of interest were ruled out by isolation of KOs of the SALK collection following genotyping with the respective forward, reverse and border primers (Alonso et al., 2003) (Supplemental Table 2). When no homozygous KO was identified (AT5G22600, AT5G22640, AT5G22650, AT5G22660, AT5G22670, AT5G22680, AT5G22710, AT5G22730), primer pairs were designed covering the full open reading frame, and amplicons obtained using *cue8* genomic DNA template submitted for Sanger sequencing (DNASeq, Medical Sciences Institute, University of Dundee, UK). When polymorphisms against the TAIR sequence occurred, amplicons for the pOCA108 WT were also sequenced (AT5G22640, AT5G22650, AT5G22660, AT5G22670, AT5G22710, AT5G22730).

### Vector construction and complementation

A Transformation-competent Artificial Chromosome (TAC (Liu et al., 2000), JatY-57L07, containing the genomic region covering genes AT5G22640 to AT5G22740, was obtained as an *E. coli* stab culture from the John Innes Centre (Norwich, UK). The TAC was introduced into *Agrobacterium* strain GV3101 by electroporation, followed by transformation of Arabidopsis *cue8* using the floral dip method (Clough and Bent, 1998). Selection of transformants utilised resistance to BASTA (Glufosinate), sprayed at 150mg/l as a mist every 3 days. Diagnostic PCR used the primers pYLTAC17-F (AATCCTGTTGCCCDCCTTG) in the vector, and 57L07-FOR-R (GTCTGAGCCAGAGCCAGAGCTTGAGG) in the insert, and produced a 607 bp amplicon. A separate TAC, JatY-76P13, covering a broader region, was employed in the same way but generated no transformants.

To produce a full-length WT cDNA, RNA was isolated (RNeasy kit, Qiagen, Manchester, UK) from pOCA108 plants, cDNA synthesised (AMV Reverse Transcriptase kit, Promega, Southampton, UK) and amplified with primers CUE8cDNA-F (CACCATGGCTAACGAAGAACTCAC) and CUE8cDNA-STOP-R (AGAGACTCAAGACACAGCAGGA) using BIO-X-ACT Long DNA polymerase (Bioline, London, UK). The 2622 bp product was directionally cloned by ligation into the pENTR/D-TOPO vector (Invitrogen/Thermo Fisher Scientific, Hemel Hempstead, UK). Digestion with EcoRV and NotI generated 2742 and 2435 bands, confirming the cloning of the full-length cDNA, and that with EcoRI and EcoRV generated 591 and 4586 bp bands, confirming the forward orientation. Sequencing confirmed the absence of errors. The TOPO vector insert was cloned into pB7WG2 vector (Karimi et al., 2002) using Gateway recombination, to produce the pB7WG2/CUE8c construct. The pB7WG2 vector includes an upstream 35S promoter. Sequencing confirmed the correct orientation and absence of errors. Transformation of pB7WG2/CUE8c used the floral dip method. Transformants were selected using BASTA. Primers cDNAgate-F (TGCCCAGCTATCTGTCACTTC) in the vector, and cDNAgate-R (CTTCCAACGTTCTGGGTCTC) in the *CUE8* sequence, generated a diagnostic 808 bp amplicon.

### *In silico* structure and expression analysis

Domain structure of the polypeptide sequence was analysed at https://www.ebi.ac.uk/interpro/protein/UniProt/. Three-dimensional predicted structure was obtained at https://alphafold.ebi.ac.uk/.

Expression of *CUE8/TIC100/EMB1211* in the Arabidopsis GeneAtlas data (Schmid et al., 2005), available at http://jsp.weigelworld.org/AtGenExpress/resources/, was compared with that of a typical photosynthetic protein, *LHCB2* (AT2G05100) and a housekeeping plastid import component, *TOC34* (AT5G05000). Gene expression correlators in relation to development, AtGenExpress tissue compendium (Schmid et al., 2005) were identified using the BioArray Resource (Toufighi et al., 2005) available at http://bar.utoronto.ca/. Coexpressors were also identified using ATTD-II (Obayashi et al., 2018).

### Chloroplast protein import assays and protein immunoblots

Chloroplasts were isolated from seedlings (Figure 3 and Figure 6) grown *in vitro* to equivalent stages of cotyledons and first leaf pair, for approximately 13 days (WT) and 17 days (mutants). Chloroplasts were isolated, examined by phase-contrast microscopy to confirm integrity, and their density quantified. Isolation and import assays using equal numbers of chloroplasts and the RBCS and RPL11 radiolabelled pre-proteins were carried out as previously described (Kubis et al., 2003). The fraction of pre-protein imported, obtained by quantifying on the same import product gel, depended on assay but was at least 10%. Extracts of total chloroplast protein were prepared and equal amounts of protein of WT and mutant were fractionated and subject to immunoblot using specific antibodies, as described (Kikuchi et al., 2013). Protein samples were denatured at 100 °C for 5 minutes except for the TIC214 (37 °C for 30 minutes) as described (Kikuchi et al., 2013). Antibody dilutions are given in the Supplemental Table 5. Quantitation of bands was carried out as described (Kubis et al., 2003) or using ImageJ software.

### RNA extraction and quantitative real-time RT-PCR analysis

Total RNA was extracted from *in vitro*-grown (under continuous light) 5-day-old wild type and 6-day-old *tic100^cue8^* seedlings. Age differences other than 24h could have resulted in spurious circadian effects. Nucleic acid extraction and quantitation, cDNA synthesis, real time-PCR amplifications, assessment of product quality and quantitation of expression in the mutant relative to that in the pOCA108 wild type was carried out as previously described (Loudya et al., 2020). Primer pairs for qRT-PCR are listed in Supplemental Table 6.

### Mutagenesis and isolation of *soh1*

The *tic100^cue8^* seeds (over 5,000) were mutagenized using 50 mM ethyl methanesulfonate for 4 hrs. About 5,000 healthy M1 *tic100^cue8^* plants (carrying heterozygous mutations) were grown as 50 pools. A putative suppressor in the M2 population from pool 18 was isolated several times and found to have a dramatic phenotype after 2 weeks on soil, which was confirmed by genotyping for the *tic100^cue8^* mutation. The protochlorophyllide and chlorophyll content in its M3 progeny seedlings further showed a clear suppression of *tic100^cue8^*. Genetic analysis of a backcross led to the conclusion of a semi-dominant suppressor mutation. Pair-wise crosses of these suppressors from pool 18 showed them to be allelic.

### Protochlorophyllide and chlorophyll content

Pigments were extracted in dimethyl formamide and quantitation was carried out by spectrophotometry or spectrofluorimetry as previously described (López-Juez et al., 1998; Vinti et al., 2005).

### Mapping by sequencing of the *soh1* mutation

The *soh1* mutation was identified by short-read mapping of a DNA pool from 150 backcrossed BC1F2 (see Supplemental Figure 5) recombinant *tic100^cue8^* phenotypes (F), as well as 100 unmutagenised *tic100^cue8^* wild types (P1) and 100 homozygous *soh1* (P2) parents. Sequencing was carried out at the Oxford Genomics Centre, Wellcome Trust Centre for Human Genetics (http://www.well.ox.ac.uk/ogc/) and mapping-by-sequencing was performed using the SHOREmap analysis package (http://bioinfo.mpipz.mpg.de/shoremap/guide.html). To narrow the region, filters were set for quality reads (>100) and indels were included to make sure the polymorphisms of Bensheim were not considered as causal mutations. To identify the semi-dominant mutation a mapping strategy was designed to first compare the polymorphisms in the *tic100^cue8^ soh1* parent (P2, test) caused by mutagenesis and that are absent in the *tic100^cue8^* parent (P1, reference) which gave list A. Secondly, the polymorphisms (induced mutations) in the backcrossed F2 *tic100^cue8^* population which are absent in P1 resulted in list B. In the last step, list A was used as a test and list B as a reference to find out the EMS-induced true SNPs.

### Gene cloning and generation of transgenic plants

Gene cloning was performed using Gateway^®^ Technology (Invitrogen). The primers used for the generation of transgenic plants and transient assays are listed in the Supplemental Table 7. The full coding sequences (CDSs) of *tic100^soh1^* and *tic100^cue8^* genes were PCR amplified from the cDNA of the respective Arabidopsis genotypes (Bensheim). The CDSs from entry and destination vectors were confirmed by sequencing (Eurofins Genomics, Constance, Germany) and transformed into the *tic100^cue8^* mutant using *Agrobacterium*-mediated transformation (floral dipping). At least 10 T1 plants resistant on BASTA plates were genotyped in each case (35S:*tic100^soh1^*and 35S:*tic100^cue8^*) and confirmed to carry the transgene.

### Subcellular localisation of TIC100 fluorescent protein fusions

To study the protein localization using YFP fluorescence, the CDSs of *TIC100, tic100^cue8^* and *tic100^soh1^* genes were PCR-amplified without the stop codon from the cDNA of their Arabidopsis parent (Bensheim genotype). The CDSs were introduced into the entry vector, sequenced and later subcloned into the plant expression vector p2GWY7 carrying a C-terminal YFP tag (Karimi et al., 2005). Protoplast isolation and transfection assays were carried out as described previously (Wu et al., 2009; Ling et al., 2012). Plasmid DNA (5 μg) was transfected to 10^5^ protoplasts (0.1 ml of protoplast suspension) isolated from healthy leaves of Arabidopsis Columbia.

The YFP fluorescence images were captured using a Leica TCS SP5 microscope as described previously (Ling et al., 2019). Images shown represent results of at least two independent experiments (at least 40 protoplasts per genotype) showing the same result.

### Lincomycin treatment and associated immunoblot

Seeds were plated and seedlings grown *in vitro* as indicated above without lincomycin. For lincomycin treatment, seeds were plated on a sterile, fine nylon mesh overlaying MS medium with 1% sucrose for 36 hours, at which time the mesh with germinating seeds was transferred to new medium containing in addition 0.5 mM lincomycin, where they continued to grow. Seedlings were harvested for transcripts’ analysis at comparable developmental stages: 5 days for wild type, 6 days for *tic100^cue8^* and *tic100^soh1^* and 7 for *tic100^soh1^ gun1*, with two additional days for protein analysis in each case. Total protein extraction using a urea/acetone powders method, incubation with a primary antibody against FtsH2 (Var2) and secondary antibody detection were as described (Loudya et al., 2021).

### Quantitation of RNA editing

Monitoring and quantitation of editing of two chloroplast mRNAs was carried out as previously described (Loudya et al., 2020).

### Statistical analyses

Averages and standard errors of the mean are indicated. Regressions, Chi-squared, Student’s t-tests (two-tailed) and ANOVA followed by Dunnett’s tests were carried out in Microsoft Excel ®, with plug-ins from Real-Statistics.com, for data using the numbers of replicates indicated for each experiment. For morphological parameters, chloroplast quantitative data, chloroplast preparations, import assays, immunoblots and gene expression assays, the number of samples represent independent biological replicates.

## Supplemental Data

**Supplemental Datasets 1 and 2.** Developmental expression co-regulators of *CUE8* identified using Arabidopsis Gene Atlas data, and expression co-regulators according to ATTED-II.

## Acknowledgments

We are grateful to Masato Nakai (Osaka University) for the 1 MDa component antibodies, Kenichi Yamaguchi (Nagasaki University) and Wataru Sakamoto (Okayama University for RPL2 and FtsH2 (Var2) antisera respectively, Elizabeth Haswell (Washington University) for the plastid-targeted DsRed line, Ian Bancroft (John Innes Centre) for the JatY clones, Korbinian Schneeberger (MPI for Plant Breeding Research) for advice on SHOREmap and John R. Bowyer (Royal Holloway) for help mapping *tic100^cue8^*. This work was funded by doctoral studentships from the UK’s BBSRC to DM, the Government of India’s Ministry of Tribal Affairs to NL and the Brunei Government Ministry of Education to SMA, by BBSRC grants BB/J009369/1, BB/K018442/1, BB/N006372/1, BB/R016984/1 and BB/R009333/1 to RPJ and BB/SB13314 to Prof. Laszlo Bogre and ELJ, and by Royal Holloway strategy funds to ELJ. Dedicated to the memory of Prof. John R. Bowyer, with whom many deeply insightful discussions were held, and Prof. Danny J. Schnell, a towering figure who pioneered our modern understanding of protein import into chloroplasts. The authors declare no competing interests.

## Author contributions

NL, DM, JB, RPJ and ELJ designed the research. DM with assistance of ELJ cloned the *CUE8* gene. NL isolated *soh1* and cloned the gene with assistance from SMA. NL and JB performed *in vitro* import assays and associated immunoblot experiments. NL performed all microscopy analysis, gene expression quantitation, cloning of fusion proteins, subcellular localisation, lincomycin-associated immunoblot and RNA editing assays. NL and ELJ performed genetics, coexpression and protein folding analyses. NL, RPJ and ELJ wrote the manuscript. PFD, RPJ and ELJ supervised the project.

---

**Supplemental Figure 1.**
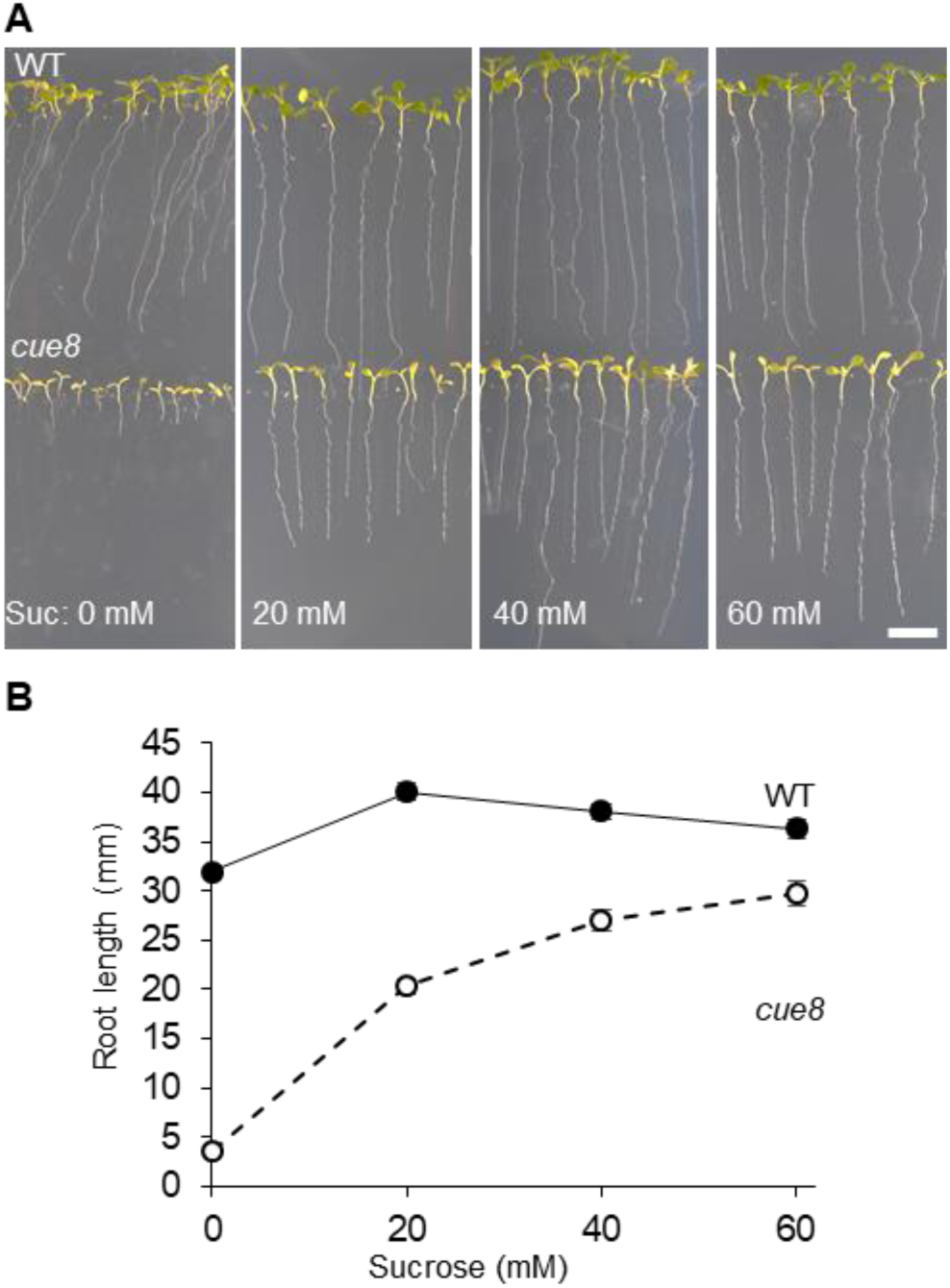
Mutation of *CUE8* delays root development, in a way which can be partly but not fully rescued by growth on sucrose. (**A**) 14-day-old seedlings of *cue8* and WT grown on vertical plates, on media containing sucrose at the concentrations indicated. Scale bar: 1 cm. (**B**) Measurement of root length of seedlings grown as above. Error bars represent s.e.m. (n ≥ 30). Values for *cue8* were significantly different to those of WT at each sucrose concentration (Student’s t-test, p<0.001). Supports Figure 1.

**Supplemental Figure 2.**
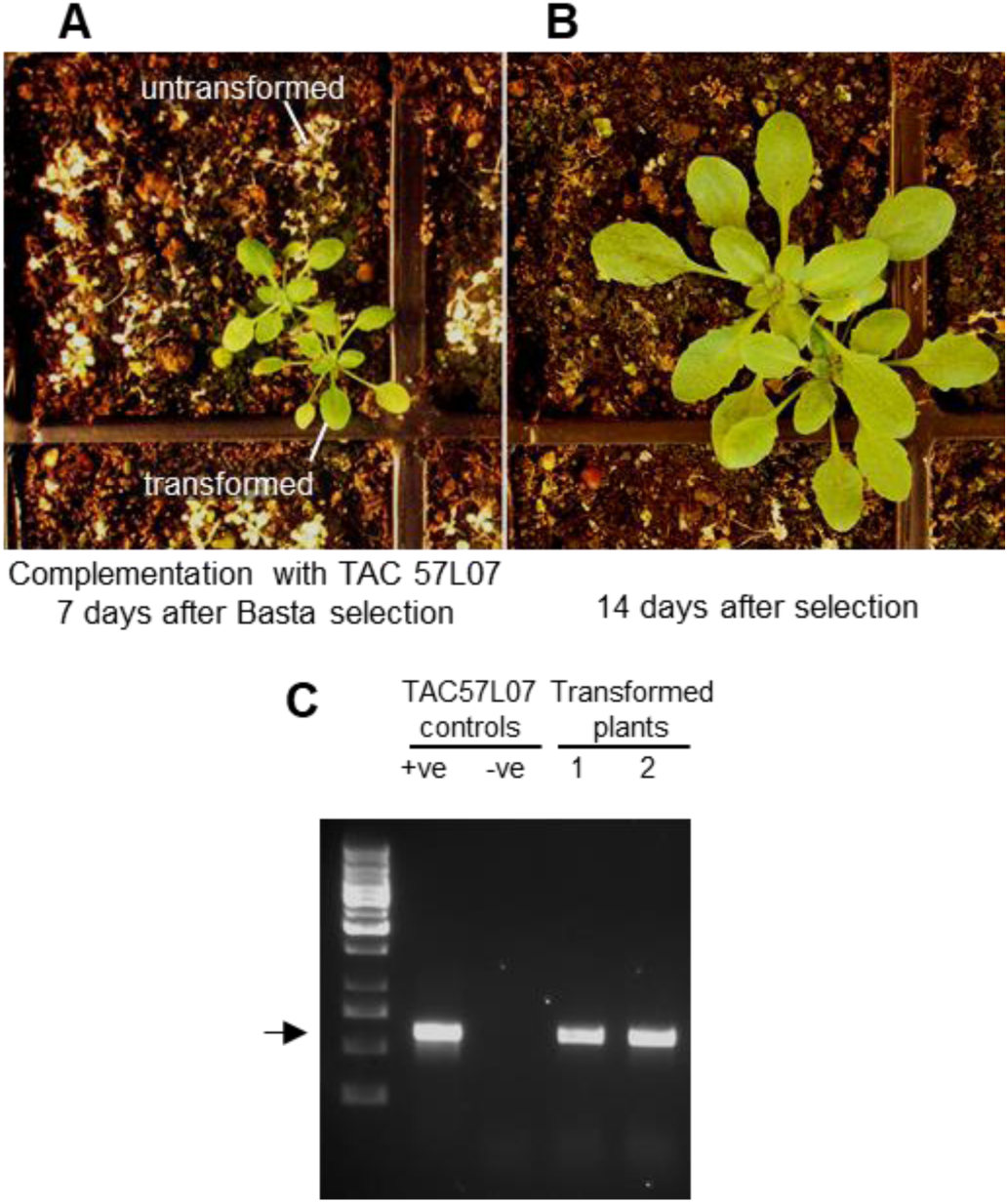
Complementation of *cue8* by genomic DNA containing *TIC100*. (**A**) pTAC JatY57L07- transformed *cue8* seedlings, sown on soil, 14 days after germination and 7 days after selection with BASTA, as described in Supplementary Materials and Methods. Note BASTA-sensitive bleached seedlings. (**B**) Same seedlings, 21 days after germination. (**C**) Diagnostic PCR amplification from complemented plants (1, 2) with one primer specific to the pYLTAC17 vector and another specific to the genomic region of TAC57L07, confirming presence of an expected 607 bp product. Positive (+ve) control, plasmid DNA harbouring the construct. Negative (-ve) control, DNA from plant prior to transformation. The second and third bands from the bottom of the left lane correspond to 500 and 750 bp respectively. Supports Figure 2.

**Supplemental Figure 3.**
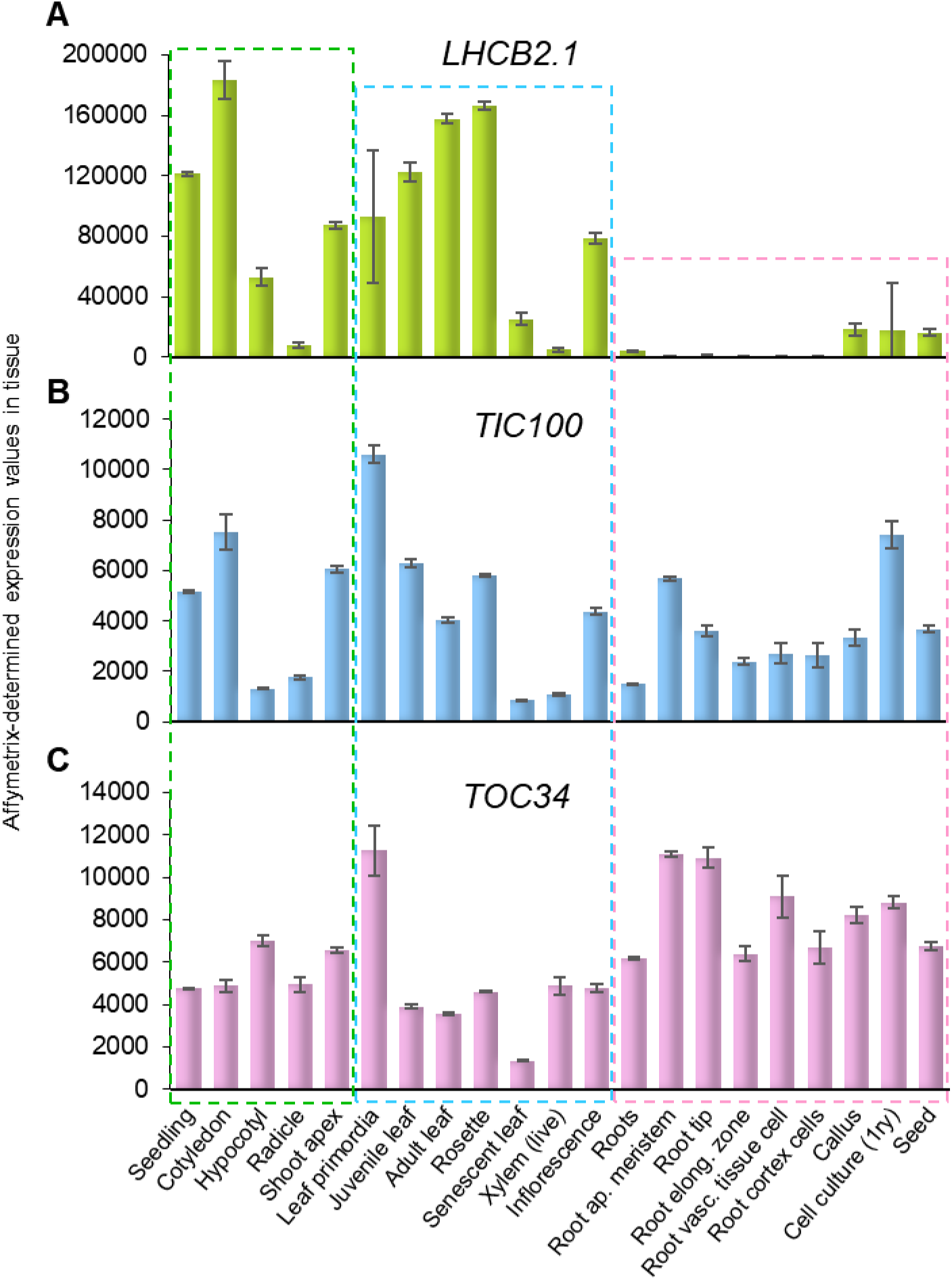
Developmental expression of *TIC100*, in relation to that of a characteristic photosynthesis-associated and a characteristic plastid housekeeping protein nucleus-encoded gene. (A) Expression of *LHCB2.1* (AT2G05100), the gene for a photosynthetic antenna polypeptide. (B) Expression of *TIC100*. (C) Expression of *TOC34* (AT2G05100), a housekeeping plastid import component gene. All developmental expression levels as identified by the Arabidopsis GeneAtlas. Supports Figures 2 and 6.

**Supplemental Figure 4.**
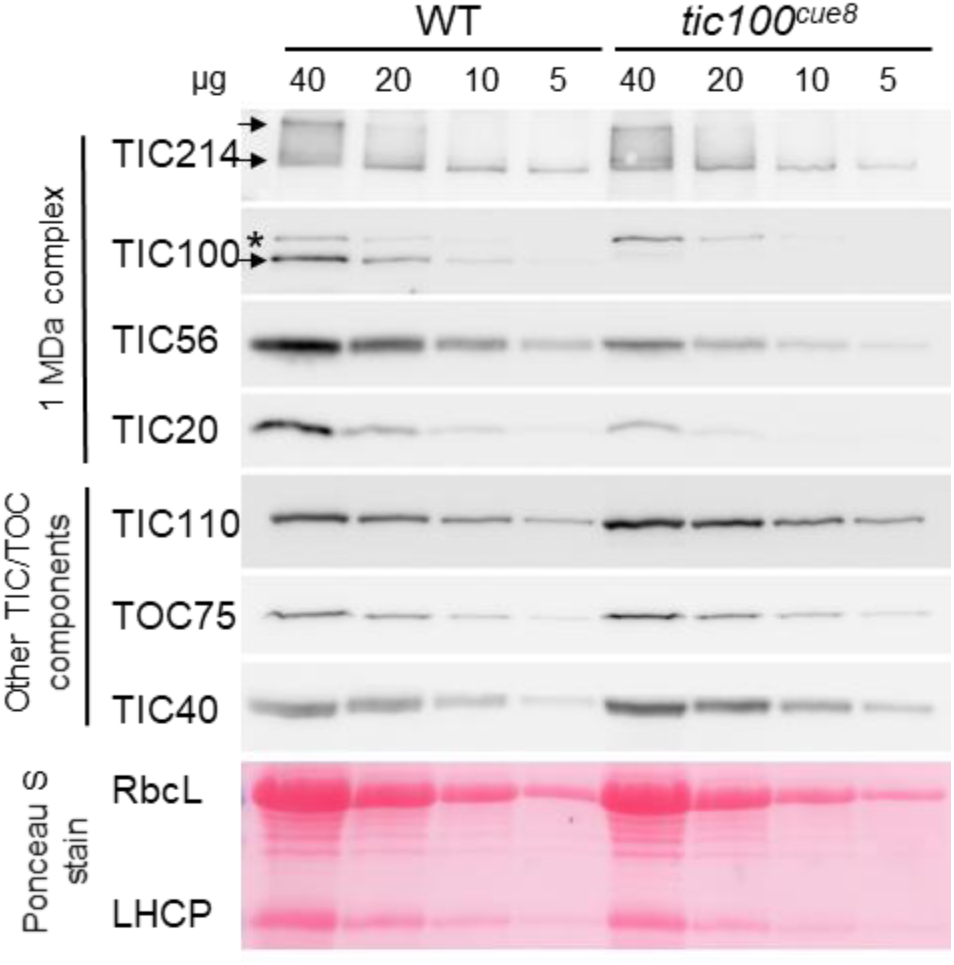
Chloroplasts of *tic100^cue8^* exhibit reduction specifically in 1MDa complex component proteins. Immunoblot analysis of total chloroplast proteins from the *tic100^cue8^* mutant and wild-type seedlings. The amount of proteins (µg) loaded is indicated above each lane. The antibodies used for the detection of components of the 1 MDa complex (TIC20, TIC56, TIC100 and TIC214) or other chloroplast envelope proteins (TOC75, TIC40 and TIC110), are indicated. In the TIC214 strip both bands, indicated by arrows, correspond to the TIC214 protein, the upper band, indicated by the top arrow, corresponding to an aggregated form of this protein caused by its large size and hydrophobic nature. The asterisk on the TIC100 strip represents a non-specific band, which serves as internal control, while the arrow corresponds to the TIC100 protein. Note the reduced amount of polypeptide components of the 1 MDa complex, in spite of the increased amount of other envelope polypeptides, including the one labelled with an asterisk in the TIC100 strip, in the *cue8* samples. The lower strip represents the Ponceau-stained total protein of one of the replica membranes. Supports Figure 3.

**Supplemental Figure 5.**
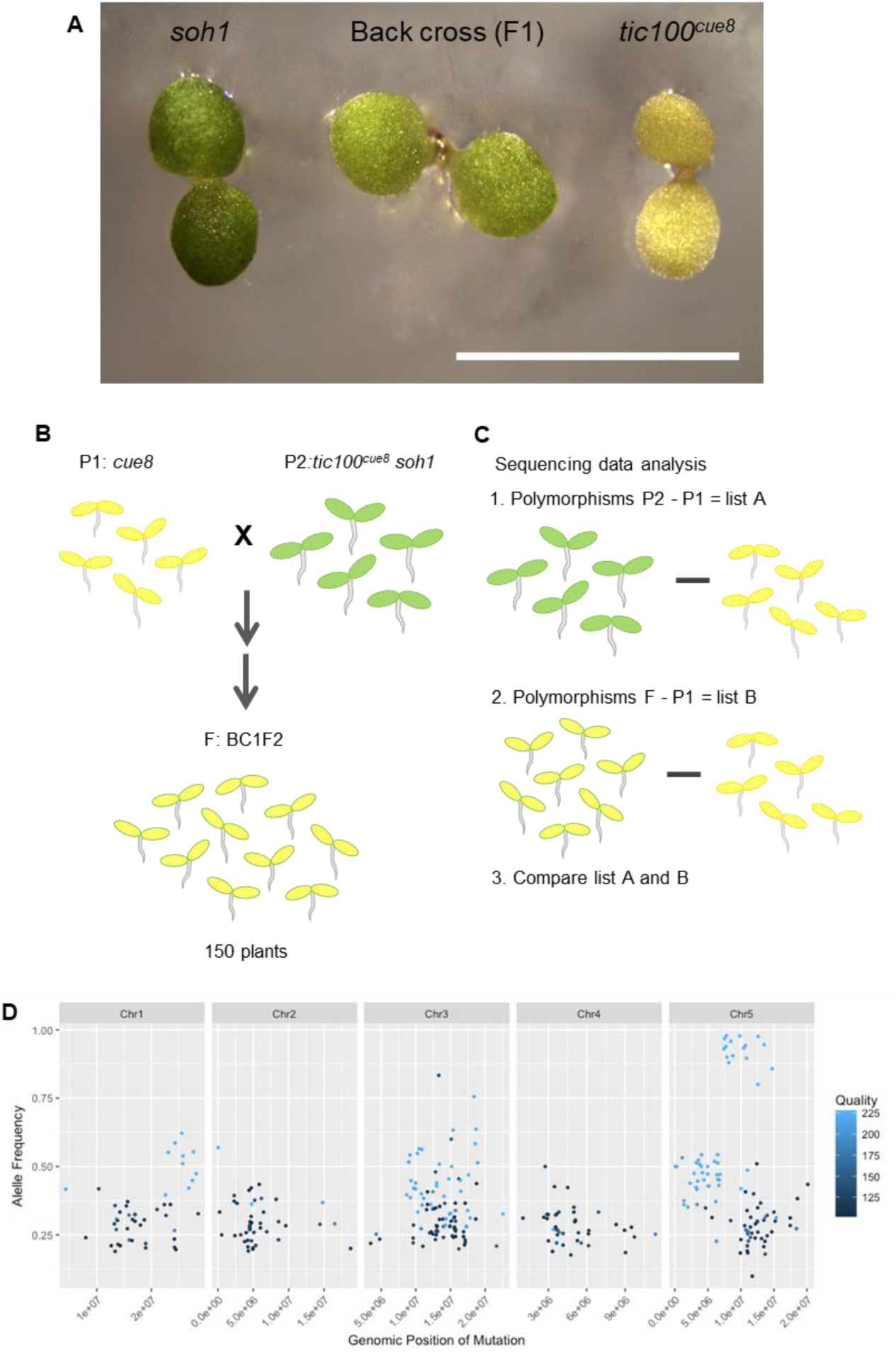
Semidominant phenotype of the *soh1* mutation, and the mapping strategy for gene identification. **(A)** Heterozygous F1 seedlings of a backcross of *soh1* (the *tic100^cue8^ soh1* double mutant) to *tic100^cue8^* are shown, together with a *tic100^cue8^* and a homozygous *soh1* seedling. Scale bar: 5 mm. Supports Figure 4. **(B)** Strategy followed for the mapping. An F2 population of phenotypically-*cue8* plants was generated from a backcross of the semidominant *soh1* mutant (P2) with its *cue8* parent (P1). This population was high-throughput sequenced in bulk, as were its two parents, before drawing the two lists of polymorphisms to compare. **(C)** Polymorphisms of both parentals (P1 and P2) and the bulked backcrossed F2 segregants population (BC1F2) relative to the Arabidopsis Genome Initiative sequence were compared as shown. **(D)** Visual output of Shoremap software demonstrating the saturation of recombinant mutant allele frequency **(B** shared with **A)** on chromosome 5 around the *TIC100* (AT5G22640) locus. **B** and **C** support Figure 5

**Supplemental Figure 6.**
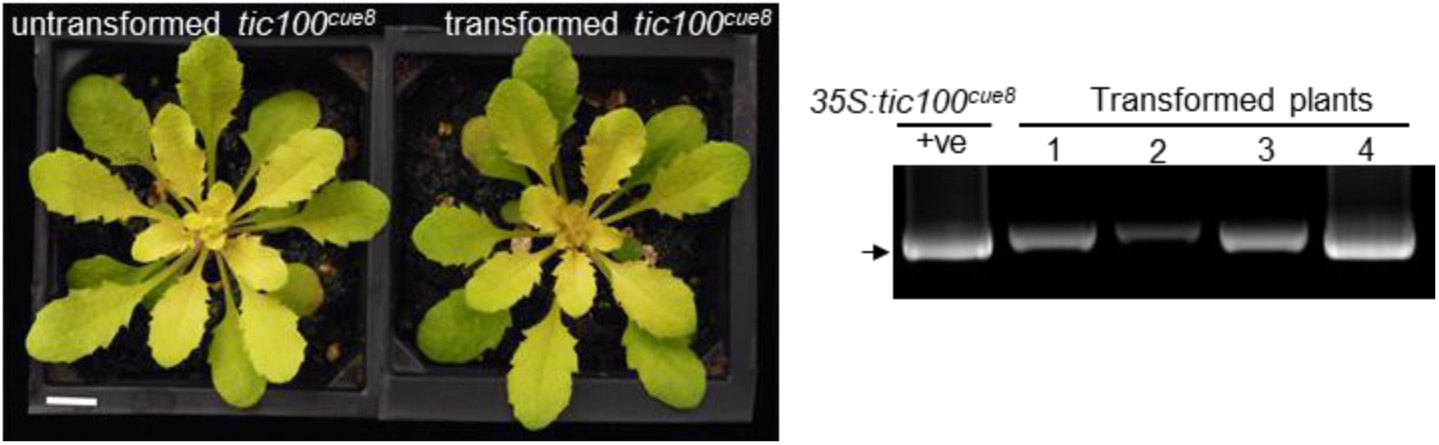
Overexpression of *tic100^cue8^* coding sequence in the *tic100^cue8^* mutant does not suppress the mutant phenotype. Failure of phenocopying of the suppressor *soh1* mutant by transformation of the single *tic100^cue8^* mutant with an over-expressed *tic100^cue8^* sequence driven by the 35S promoter (as seen in 4 independent T1 plants). Plants shown at 40 days of age. Scale bar: 1cm. Gel on the right confirms the genotype of the transformed plants. “+ve”: positive genotyping control (bacterial plasmid). Supports Figure 5E.

**Supplemental Figure 7.**
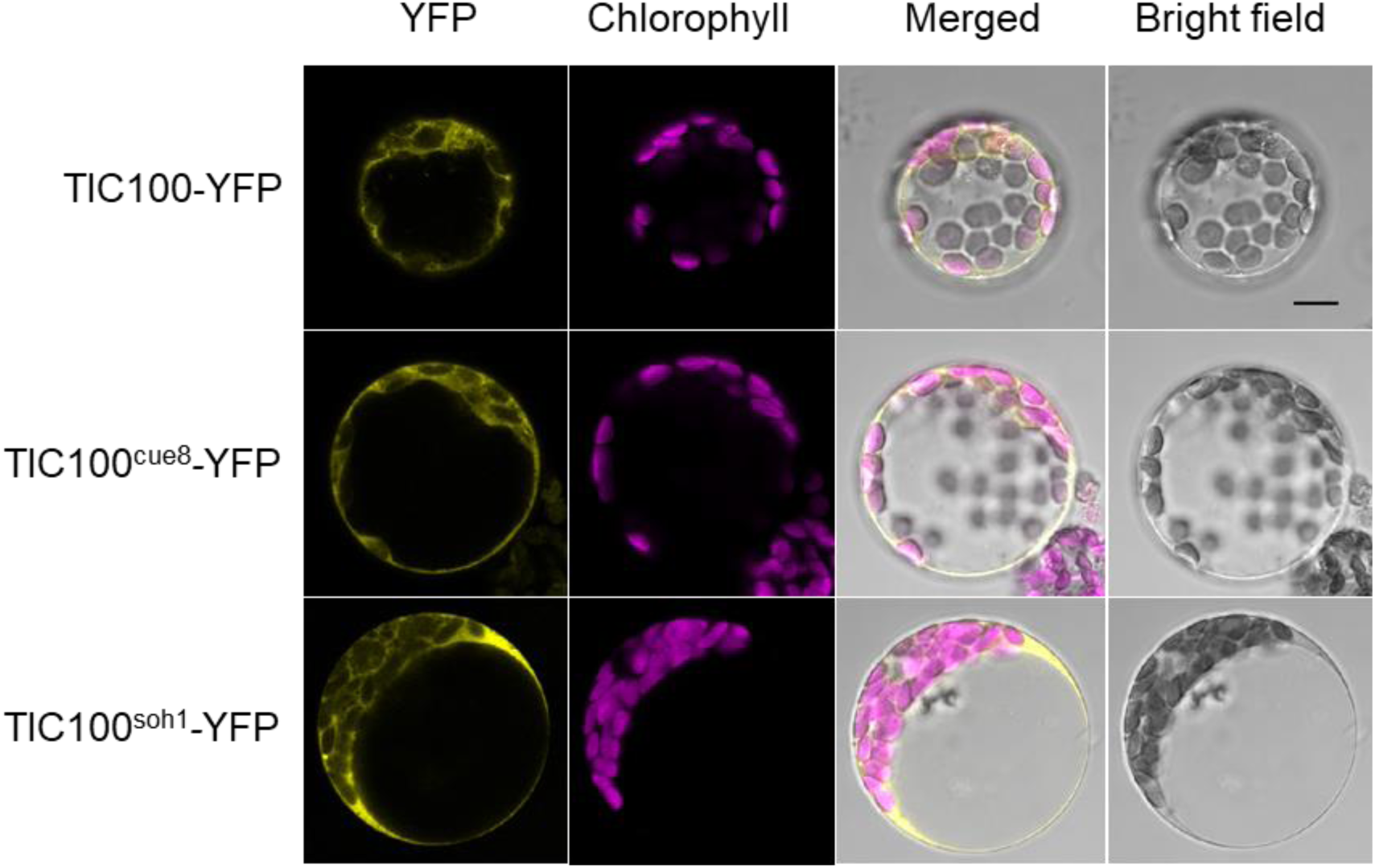
Localisation of the TIC100 protein, in its wild type, TIC100^cue8^ and double-mutated TIC100^soh1^ forms, to the cytoplasm and the chloroplast periphery of transformed, over-expressing intact protoplasts. Wild-type protoplasts were transfected with constructs encoding wild-type and mutant forms of TIC100, each one tagged with a C-terminal YFP tag, before observation of the fusion protein using confocal microscopy. Scale bar: 10 µm. Supports Figure 5F.

**Supplemental Figure 8.**
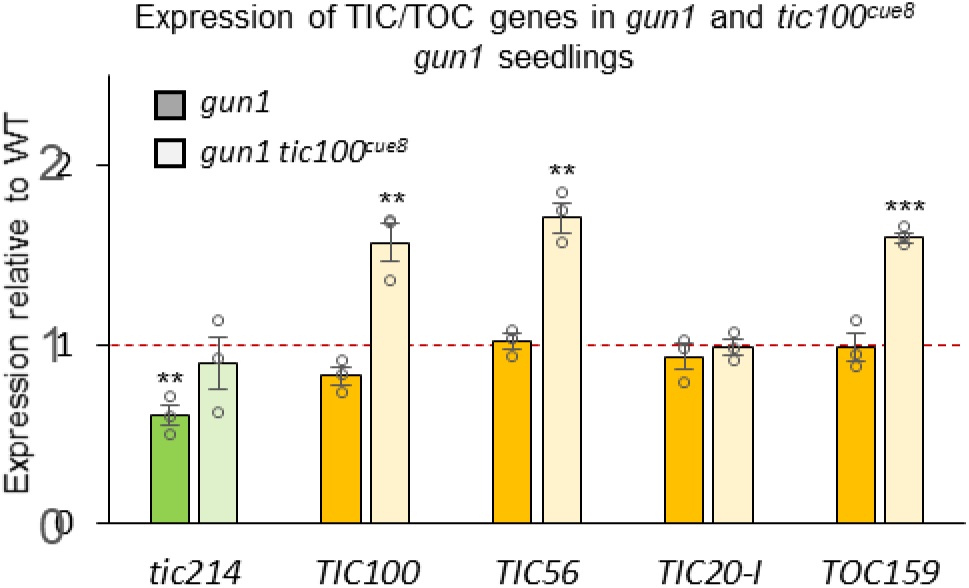
The increase in expression of genes for components of the 1MDa complex observed in *tic100^cue8^* (Fig. 7) is only partially dependent on the action of GUN1, and can be observed to some extent even in GUN1 absence. Expression, measured by quantitative real-time RT-PCR, of *TIC*/*TOC* genes in *tic100^cue8^ gun1* seedlings, measured relative to expression in wild-type seedlings and compared to expression in *gun1*. Note *tic214* is chloroplast-encoded. The presented values are means, and the error bars show s.e.m. of three RNA samples (biological replicates), each with two technical replicates. Asterisks represent significance of difference between mutant and WT (as indicated for Figure 7D, 2-tailed Student’s t-test). Dotted lines represent expression in WT. Supports Figure 7.

**Supplemental Table 1.**
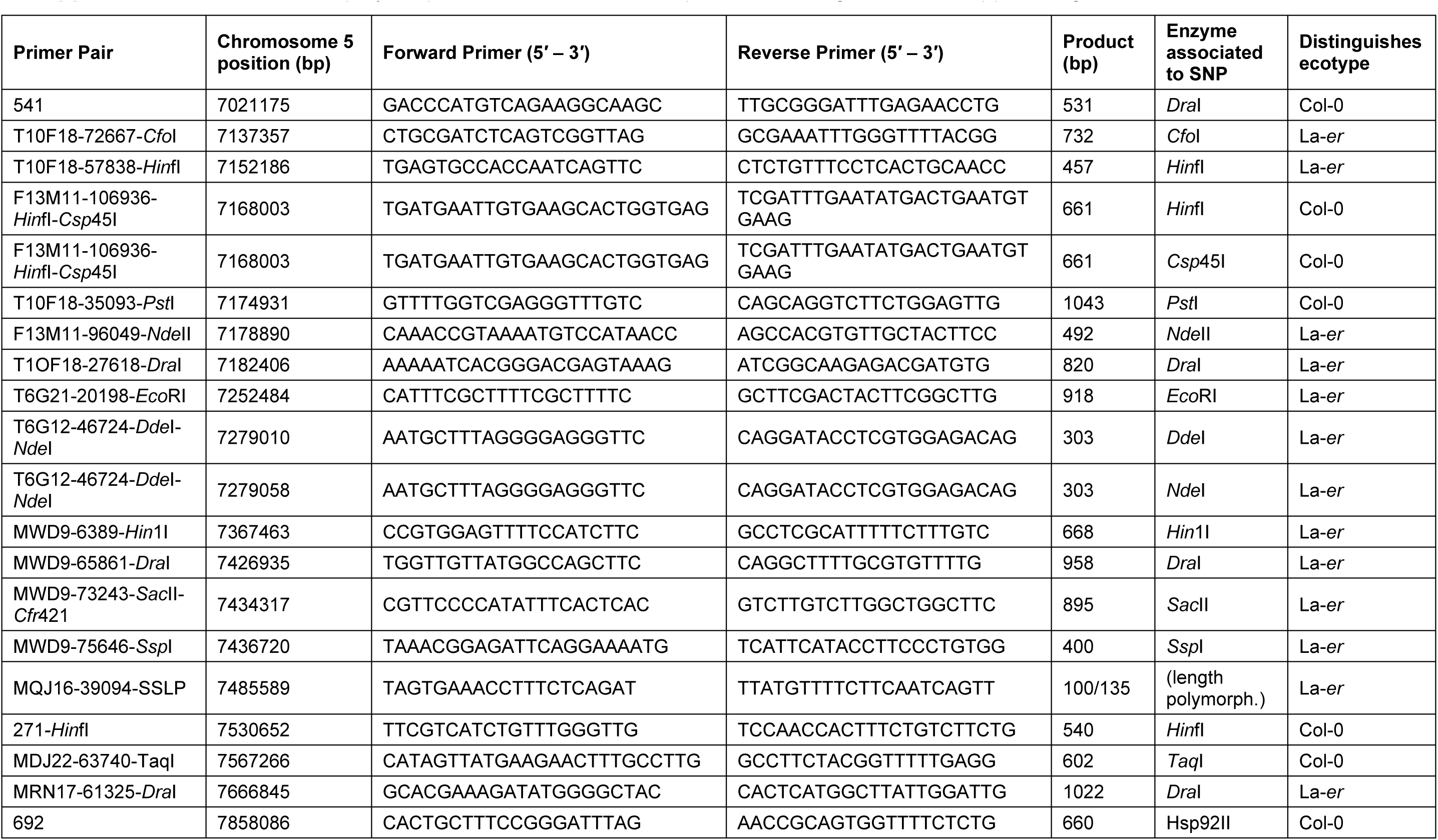
List of polymorphic markers used for map-based cloning of *CUE8.* Supports Figure 2

**Supplemental Table 2.** Analysis of the genomic region containing the *CUE8* gene, and strategies used to rule out alternatives. Supports Figure 2.

| Accession | Gene product | Mutation ruled out by complementing TAC? | T-DNA line | Mutation ruled out by T-DNA phenotype? | Mutation ruled out by sequencing? | *CUE8* |
| --- | --- | --- | --- | --- | --- | --- |
| AT5G22555 | Expressed protein | No |  |  | ✓ |  |
| AT5G22560 | Hypothetical protein | No |  |  | ✓ |  |
| AT5G22570 | Transcription Factor | No |  |  | ✓ |  |
| AT5G22580 | Expressed protein | No |  |  | ✓ |  |
| AT5G22590 | Hypothetical protein | No | SALK_144064 | ✓ | ✓ |  |
| AT5G22600 | Expressed protein | No | SALK_093885 | No T-DNA | ✓ |  |
| AT5G22610 | F-box family protein | No | SALK_117573 | ✓ |  |  |
| AT5G22620 | Phosphoglycer-ate mutase | No | SALK_012577 | ✓ |  |  |
| AT5G22630 | Prephenate dehydratase | No | SALK_028611 | ✓ |  |  |
| AT5G22640 | TIC100 | Complemented by<br>JatY-57L07 | SALK_138825 | Embryo/<br>seedling lethal | G2087A<br>(gly366arg) | ✓ |
| AT5G22650 | Histone deacetylases |  |  |  | ✓ |  |
| AT5G22660 | F-box family protein |  |  |  | ✓ |  |
| AT5G22670 | F-box family protein |  |  |  | ✓ |  |
| AT5G22680 | Hypothetical protein |  |  |  | ✓ |  |
| AT5G22690 | Disease resistance protein |  | SALK_039393 | ✓ |  |  |
| AT5G22700 | F-box family protein |  | SALK_009942 | ✓ |  |  |
| AT5G22720 | F-box family protein |  | SALK_002785 | ✓ |  |  |
| AT5G22730 | F-box family protein |  |  |  | ✓ |  |
| AT5G22740 | Cellulose synthase |  | SALK_149092 | ✓ |  |  |

**Supplemental Table 3.**
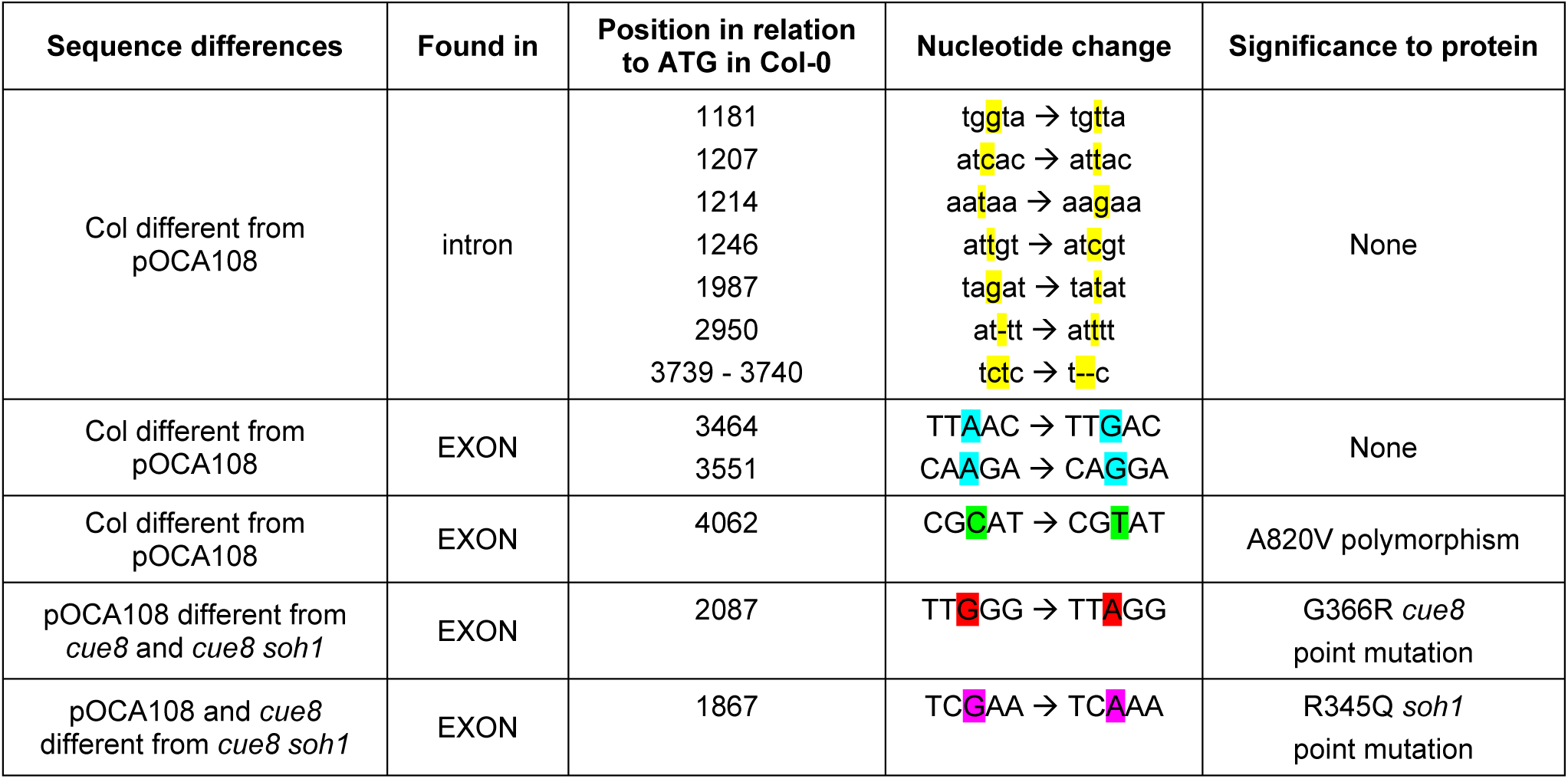
Polymorphisms in the *TIC100* sequence between the different genotypes and mutants. Supports Figure 2. Exon (upper case), intron (low case). In addition to the G366R mutation in *cue8* and the additional R345Q mutation in *cue8 soh1*, ten other polymorphisms were observed against the Arabidopsis Genome Initiative (Col) sequence. Red highlight: mutation in *cue8*. Purple: second mutation in *cue8 soh1*. Yellow, blue and green: polymorphisms between *cue8*/*cue8 soh1*/pOCA108 and Col within introns (yellow) and exons (blue, green). Two polymorphisms in exons confer silent changes (blue), while the third (green) results in a single, conservative amino acid substitution (A820V), distinguishing pOCA108 and Col.

**Supplemental Table 4.**
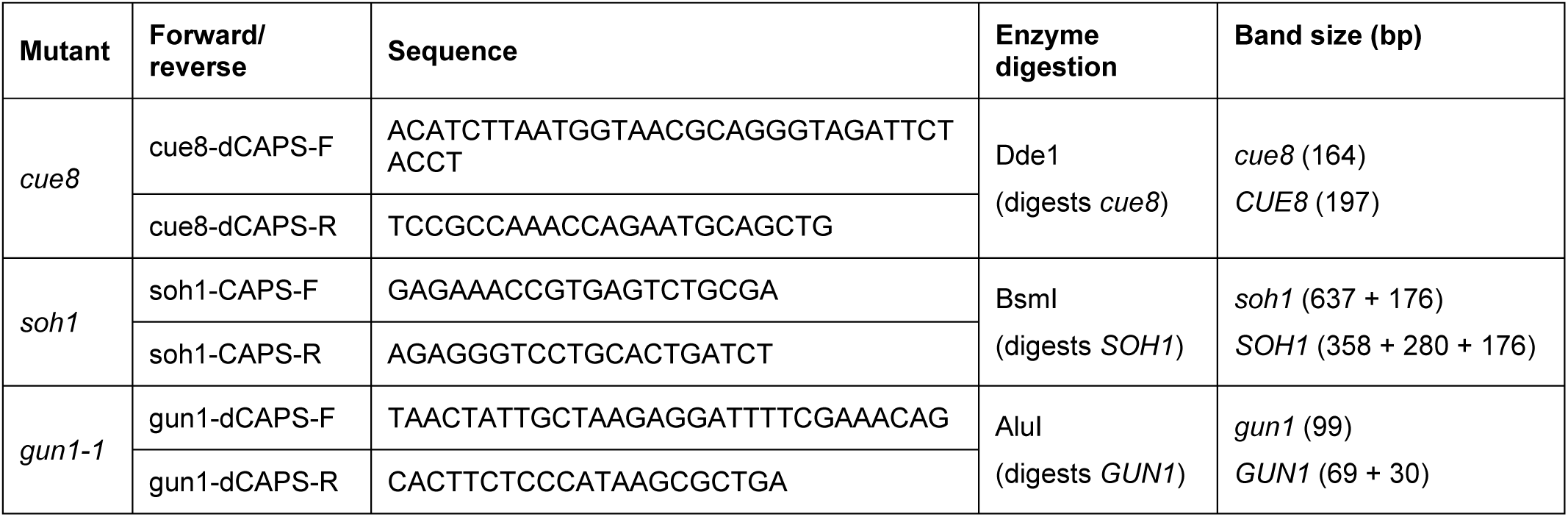
Primers used for genotyping the point mutants by dCAPS /CAPS.

**Supplemental Table 5.** Antibody dilutions used in immunoblotting.

| Antibody | Dilution | Source |
| --- | --- | --- |
| Anti-TIC214 | 1:5,000 | Kikuchi et al., 2013 |
| Anti-TIC100 | 1:5,000 |  |
| Anti-TIC56 | 1:5,000 |  |
| Anti-TIC20 | 1:50 |  |
| Anti-TIC110 | 1:5,000 | Ling et al., 2012 |
| Anti-TIC40 | 1:1,000,000 |  |
| Anti-TOC75 | 1:1,000 |  |
| Anti-HSP70 | 1:5,000 |  |
| Anti-RPL2 | 1:5,000 | Subramanian lab, Berlin |
| Anti-FtsH2 (Anti-Var2) | 1:5,000 | Sakamoto lab, Okayama |

**Supplemental Table 6.**
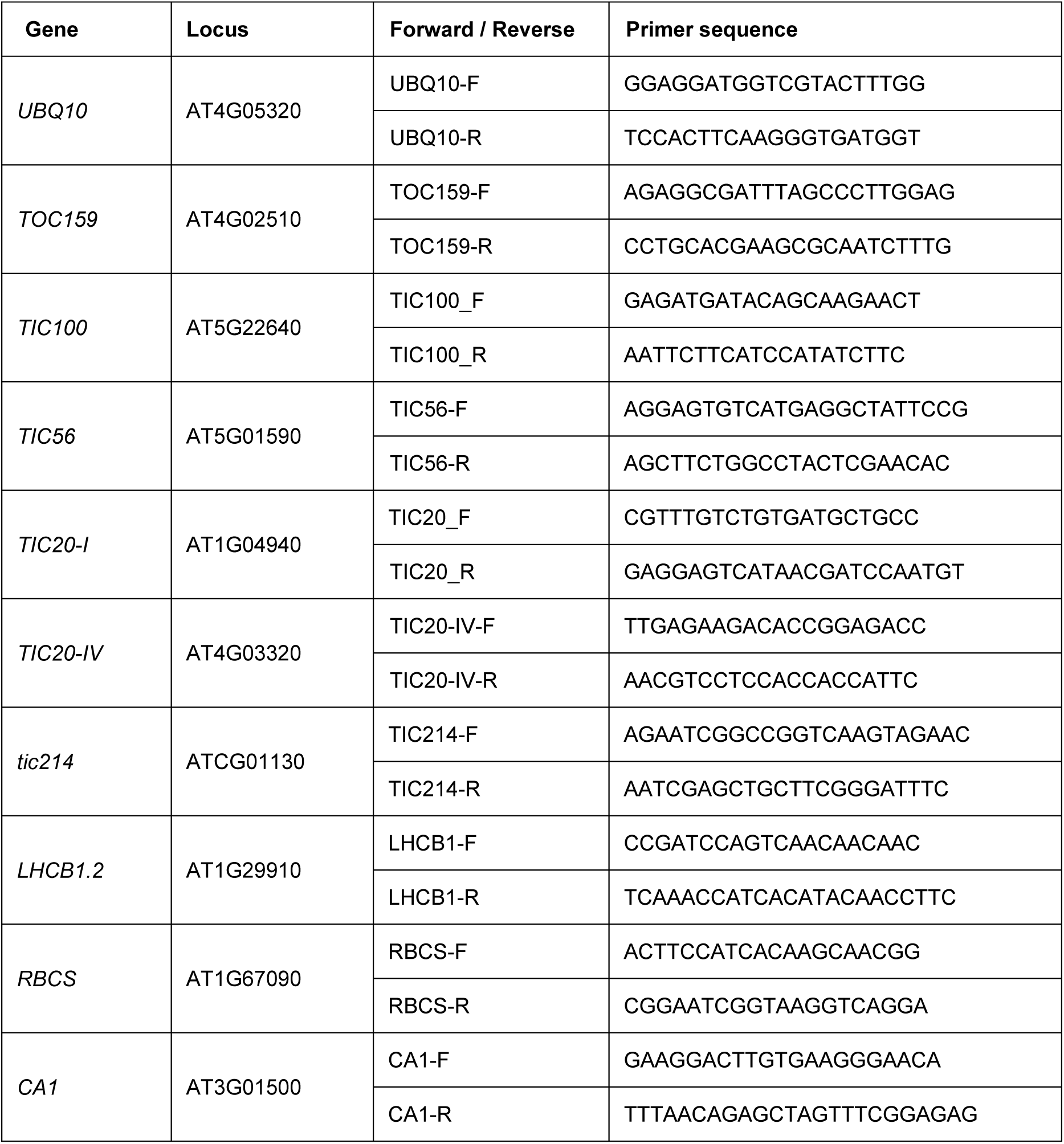
List of primers used for quantitative real-time RT-PCR.

**Supplemental Table 7.**
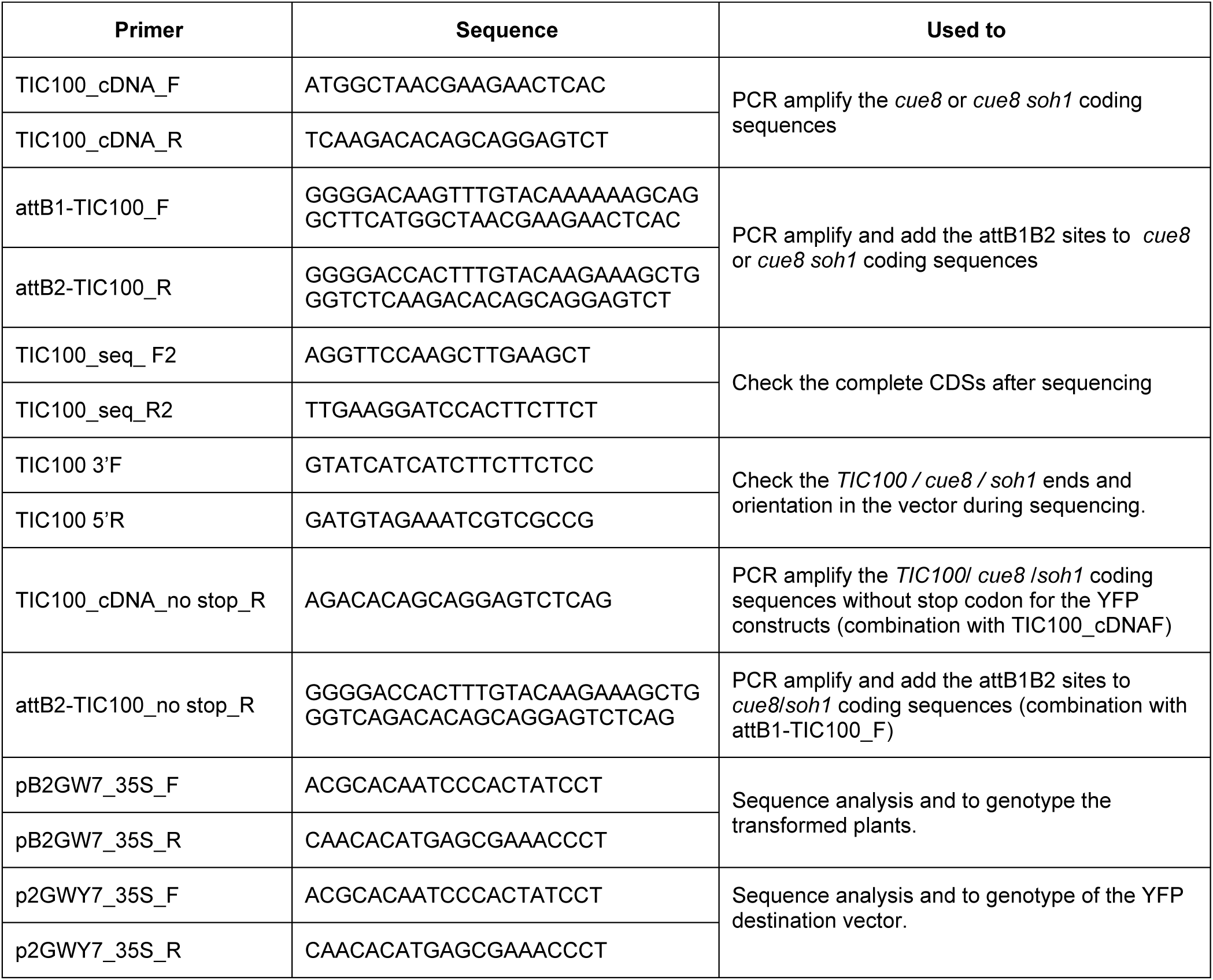
Primers used for gene cloning and transgenic approaches.

## Parsed Citations

Alonso, J.M., Stepanova, A.N., Leisse, T.J., Kim, C.J., Chen, H.M., Shinn, P., Stevenson, D.K., Zimmerman, J., Barajas, P., Cheuk, R., Gadrinab, C., Heller, C., Jeske, A., Koesema, E., Meyers, C.C., Parker, H., Prednis, L., Ansari, Y., Choy, N., Deen, H., Geralt, M., Hazari, N., Hom, E., Karnes, M., Mulholland, C., Ndubaku, R., Schmidt, I., Guzman, P., Aguilar-Henonin, L., Schmid, M., Weigel, D., Carter, D.E., Marchand, T., Risseeuw, E., Brogden, D., Zeko, A., Crosby, W.L., Berry, C.C., and Ecker, J.R. (2003). Genome-wide Insertional mutagenesis of Arabidopsis thaliana. Science 301, 653–657.

Arsovski, A.A., Galstyan, A., Guseman, J.M., and Nemhauser, J.L. (2012). Photomorphogenesis. The Arabidopsis book / American Society of Plant Biologists 10, e0147.

Bolter, B., and Soll, J. (2016). Once upon a Time -Chloroplast Protein Import Research from Infancy to Future Challenges. Mol Plant 9, 798–812.

Bolter, B., and Soll, J. (2017). Ycf1/Tic214 Is Not Essential for the Accumulation of Plastid Proteins. Mol Plant 10, 219–221.

Borner, T., Aleynikova, A.Y., Zubo, Y.O., and Kusnetsov, V.V. (2015). Chloroplast RNApolymerases: Role in chloroplast biogenesis. Bba-Bioenergetics 1847, 761–769.

Cackett, L., Luginbuehl, L.H., Schreier, T.B., Lopez-Juez, E., and Hibberd, J.M. (2021). Chloroplast development in green plant tissues: the interplay between light, hormone, and transcriptional regulation. New Phytol.

Chen, L.j., and Li, H.m. (2017). Stable megadalton TOC–TIC supercomplexes as major mediators of protein import into chloroplasts. The Plant Journal 92, 178–188.

Chen, X.J., Smith, M.D., Fitzpatrick, L., and Schnell, D.J. (2002). In vivo analysis of the role of atTic20 in protein import into chloroplasts. Plant Cell 14, 641–654.

Chotewutmontri, P., and Barkan, A. (2016). Dynamics of Chloroplast Translation during Chloroplast Differentiation in Maize. Plos Genet 12.

Chu, C.C., Swamy, K., and Li, H.M. (2020). Tissue-Specific Regulation of Plastid Protein Import via Transit-Peptide Motifs. Plant Cell 32, 1204–1217.

Clough, S.J., and Bent, A.F. (1998). Floral dip: a simplified method for Agrobacterium-mediated transformation of Arabidopsis thaliana. Plant J 16, 735–743.

Dahlin, C., and Cline, K. (1991). Developmental regulation of the plastid protein import apparatus. The Plant Cell 3, 1131–1140.

de Vries, J., Sousa, F.L., Bolter, B., Soll, J., and Gould, S.B. (2015). Ycf1: AGreen Tic? Plant Cell 27, 1827–1833.

Demarsy, E., Lakshmanan, A.M., and Kessler, F. (2014). Border control: selectivity of chloroplast protein import and regulation at the TOC-complex. Frontiers in plant science 5, 483.

Haswell, E.S., and Meyerowitz, E.M. (2006). MscS-like proteins control plastid size and shape in Arabidopsis thaliana. Curr Biol 16, 1–11.

Heins, L., Mehrle, A., Hemmler, R., Wagner, R., Kuchler, M., Hormann, F., Sveshnikov, D., and Soll, J. (2002). The preprotein conducting channel at the inner envelope membrane of plastids. EMBO Journal 21, 2616–2625.

Inaba, T., Li, M., Alvarez-Huerta, M., Kessler, F., and Schnell, D.J. (2003). atTic110 functions as a scaffold for coordinating the stromal events of protein import into chloroplasts. Journal of Biological Chemistry 278, 38617–38627.

Ivanova, Y., Smith, M.D., Chen, K., and Schnell, D.J. (2004). Members of the Toc159 import receptor family represent distinct pathways for protein targeting to plastids. Molecular Biology of the Cell 15, 3379–3392.

Jarvis, P. (2008). Targeting of nucleus-encoded proteins to chloroplasts in plants. New Phytol 179, 257–285.

Jarvis, P., and López-Juez, E. (2013). Biogenesis and homeostasis of chloroplasts and other plastids. Nature reviews. Molecular cell biology 14, 787–802.

Jumper, J., Evans, R., Pritzel, A., Green, T., Figurnov, M., Ronneberger, O., Tunyasuvunakool, K., Bates, R., Zidek, A., Potapenko, A., Bridgland, A., Meyer, C., Kohl, S.A.A., Ballard, A.J., Cowie, A., Romera-Paredes, B., Nikolov, S., Jain, R., Adler, J., Back, T., Petersen, S., Reiman, D., Clancy, E., Zielinski, M., Steinegger, M., Pacholska, M., Berghammer, T., Bodenstein, S., Silver, D., Vinyals, O., Senior, A.W., Kavukcuoglu, K., Kohli, P., and Hassabis, D. (2021). Highly accurate protein structure prediction with AlphaFold. Nature.

Kakizaki, T., Yazu, F., Nakayama, K., Ito-Inaba, Y., and Inaba, T. (2012). Plastid signalling under multiple conditions is accompanied by a common defect in RNAediting in plastids. J Exp Bot 63, 251–260.

Karimi, M., Inze, D., and Depicker, A. (2002). GATEWAY vectors for Agrobacterium-mediated plant transformation. Trends Plant Sci 7, 193–195.

Karimi, M., De Meyer, B., and Hilson, P. (2005). Modular cloning in plant cells. Trends Plant Sci 10, 103–105.

Kasmati, A.R., Topel, M., Patel, R., Murtaza, G., and Jarvis, P. (2011). Molecular and genetic analyses of Tic20 homologues in Arabidopsis thaliana chloroplasts. Plant J 66, 877–889.

Kessler, F., and Blobel, G. (1996). Interaction of the protein import and folding machineries in the chloroplast. P Natl Acad Sci USA 93, 7684–7689.

Kikuchi, S., Bedard, J., Hirano, M., Hirabayashi, Y., Oishi, M., Imai, M., Takase, M., Ide, T., and Nakai, M. (2013). Uncovering the protein translocon at the chloroplast inner envelope membrane. Science 339, 571–574.

Kikuchi, S., Asakura, Y., Imai, M., Nakahira, Y., Kotani, Y., Hashiguchi, Y., Nakai, Y., Takafuji, K., Bedard, J., Hirabayashi-Ishioka, Y., Mori, H., Shiina, T., and Nakai, M. (2018). AYcf2-FtsHi Heteromeric AAA-ATPase Complex Is Required for Chloroplast Protein Import. Plant Cell 30, 2677–2703.

Klepikova, A.V., Kasianov, A.S., Gerasimov, E.S., Logacheva, M.D., and Penin, A.A. (2016). Ahigh resolution map of the Arabidopsis thaliana developmental transcriptome based on RNA-seq profiling. Plant J 88, 1058–1070.

Kohler, D., Helm, S., Agne, B., and Baginsky, S. (2016). Importance of Translocon Subunit Tic56 for rRNAProcessing and Chloroplast Ribosome Assembly. Plant Physiol 172, 2429–2444.

Kohler, D., Montandon, C., Hause, G., Majovsky, P., Kessler, F., Baginsky, S., and Agne, B. (2015). Characterization of Chloroplast Protein Import without Tic56, a Component of the 1-Megadalton Translocon at the Inner Envelope Membrane of Chloroplasts. Plant Physiol 167, 972–990.

Koussevitzky, S., Nott, A., Mockler, T.C., Hong, F., Sachetto-Martins, G., Surpin, M., Lim, I.J., Mittler, R., and Chory, J. (2007). Signals fromchloroplasts converge to regulate nuclear gene expression. Science 316, 715–719.

Kovacs-Bogdan, E., Benz, J.P., Soll, J., and Bolter, B. (2011). Tic20 forms a channel independent of Tic110 in chloroplasts. Bmc Plant Biol 11, 133.

Kubis, S., Baldwin, A., Patel, R., Razzaq, A., Dupree, P., Lilley, K., Kurth, J., Leister, D., and Jarvis, P. (2003). The Arabidopsis ppi1 mutant is specifically defective in the expression, chloroplast import, and accumulation of photosynthetic proteins. Plant Cell 15, 1859–1871.

Kubis, S., Patel, R., Combe, J., Bedard, J., Kovacheva, S., Lilley, K., Biehl, A., Leister, D., Rios, G., Koncz, C., and Jarvis, P. (2004). Functional specialization amongst the Arabidopsis Toc159 family of chloroplast protein import receptors. Plant Cell 16, 2059–2077.

Li, H.M., Culligan, K., Dixon, R.A., and Chory, J. (1995). Cue1 - a Mesophyll Cell-Specific Positive Regulator of Light-Controlled Gene-Expression in Arabidopsis. Plant Cell 7, 1599–1610.

Liang, Q.J., Lu, X.D., Jiang, L., Wang, C.Y., Fan, Y.L., and Zhang, C.Y. (2010). EMB1211 is required for normal embryo development and influences chloroplast biogenesis in Arabidopsis. Physiol Plantarum 140, 380–394.

Ling, Q., Huang, W., Baldwin, A., and Jarvis, P. (2012). Chloroplast biogenesis is regulated by direct action of the ubiquitin-proteasome system. Science 338, 655–659.

Ling, Q., Broad, W., Trosch, R., Topel, M., Demiral Sert, T., Lymperopoulos, P., Baldwin, A., and Jarvis, R.P. (2019). Ubiquitin-dependent chloroplast-associated protein degradation in plants. Science 363, 836–848.

Liu, Y.G., Nagaki, K., Fujita, M., Kawaura, K., Uozumi, M., and Ogihara, Y. (2000). Development of an efficient maintenance and screening system for large-insert genomic DNAlibraries of hexaploid wheat in a transformation-competent artificial chromosome (TAC) vector. Plant J 23, 687–695.

López-Juez, E., Jarvis, R.P., Takeuchi, A., Page, A.M., and Chory, J. (1998). New Arabidopsis cue mutants suggest a close connection between plastid- and phytochrome regulation of nuclear gene expression. Plant Physiol 118, 803–815.

Loudya, N., Okunola, T., He, J., Jarvis, P., and López-Juez, E. (2020). Retrograde signalling in a virescent mutant triggers an anterograde delay of chloroplast biogenesis that requires GUN1 and is essential for survival. Philosophical transactions of the Royal Society of London. Series B, Biological sciences 375, 20190400.

Loudya, N., Mishra, P., Takahagi, K., Uehara-Yamaguchi, Y., Inoue, K., Bogre, L., Mochida, K., and Lopez-Juez, E. (2021). Cellular and transcriptomic analyses reveal two-staged chloroplast biogenesis underpinning photosynthesis build-up in the wheat leaf. Genome Biol 22.

Lubeck, J., Soll, J., Akita, M., Nielsen, E., and Keegstra, K. (1996). Topology of IEP110, a component of the chloroplastic protein import machinery present in the inner envelope membrane. EMBO Journal 15, 4230–4238.

Nakai, M. (2015a). The TIC complex uncovered: The alternative view on the molecular mechanism of protein translocation across the inner envelope membrane of chloroplasts. Bba-Bioenergetics 1847, 957–967.

Nakai, M. (2015b). YCF1: AGreen TIC: Response to the de Vries et al. Commentary. Plant Cell 27, 1834–1838.

Nakai, M. (2018). New Perspectives on Chloroplast Protein Import. Plant Cell Physiol 59, 1111–1119.

Nakai, M. (2020). Reply: The Revised Model for Chloroplast Protein Import. Plant Cell 32, 543–546.

Obayashi, T., Aoki, Y., Tadaka, S., Kagaya, Y., and Kinoshita, K. (2018). ATTED-II in 2018: APlant Coexpression Database Based on Investigation of the Statistical Property of the Mutual Rank Index. Plant Cell Physiol 59, 440.

Ramundo, S., Asakura, Y., Salome, P.A., Strenkert, D., Boone, M., Mackinder, L.C.M., Takafuji, K., Dinc, E., Rahire, M., Crevecoeur, M., Magneschi, L., Schaad, O., Hippler, M., Jonikas, M.C., Merchant, S., Nakai, M., Rochaix, J.D., and Walter, P. (2020). Coexpressed subunits of dual genetic origin define a conserved supercomplex mediating essential protein import into chloroplasts. P Natl Acad Sci USA 117, 32739–32749.

Richardson, L.G.L., and Schnell, D.J. (2020). Origins, function, and regulation of the TOC-TIC general protein import machinery of plastids. J Exp Bot 71, 1226–1238.

Richardson, L.G.L., Small, E.L., Inoue, H., and Schnell, D.J. (2018). Molecular Topology of the Transit Peptide during Chloroplast Protein Import. Plant Cell 30, 1789–1806.

Schafer, P., Helm, S., Kohler, D., Agne, B., and Baginsky, S. (2019). Consequences of impaired 1-MDa TIC complex assembly for the abundance and composition of chloroplast high-molecular mass protein complexes. PLoS One 14, e0213364.

Schmid, M., Davison, T.S., Henz, S.R., Pape, U.J., Demar, M., Vingron, M., Scholkopf, B., Weigel, D., and Lohmann, J.U. (2005). A gene expression map of Arabidopsis thaliana development. Nat Genet 37, 501–506.

Schneeberger, K., Ossowski, S., Lanz, C., Juul, T., Petersen, A.H., Nielsen, K.L., Jørgensen, J.-E., Weigel, D., and Andersen,S.U.J.N.m. (2009). SHOREmap: simultaneous mapping and mutation identification by deep sequencing. Nat Methods 6, 550.

Sjuts, I., Soll, J., and Bolter, B. (2017). Import of Soluble Proteins into Chloroplasts and Potential Regulatory Mechanisms. Frontiers in plant science 8, 168.

Susek, R.E., Ausubel, F.M., and Chory, J.J.C. (1993). Signal transduction mutants of Arabidopsis uncouple nuclear CAB and RBCS gene expression fromchloroplast development. Cell 74, 787–799.

Tadini, L., Pesaresi, P., Kleine, T., Rossi, F., Guljamow, A., Sommer, F., Mühlhaus, T., Schroda, M., Masiero, S., and Pribil, M.J.P.p. (2016). GUN1 controls accumulation of the plastid ribosomal protein S1 at the protein level and interacts with proteins involved in plastid protein homeostasis. Plant Physiol, pp. 02033.02015.

Tadini, L., Peracchio, C., Trotta, A., Colombo, M., Mancini, I., Jeran, N., Costa, A., Faoro, F., Marsoni, M., Vannini, C., Aro, E.M., and Pesaresi, P. (2020). GUN1 influences the accumulation of NEP-dependent transcripts and chloroplast protein import in Arabidopsis cotyledons upon perturbation of chloroplast protein homeostasis. Plant J 101, 1198–1220.

Takeshima, H., Komazaki, S., Nishi, M., Iino, M., and Kangawa, K. (2000). Junctophilins: a novel family of junctional membrane complex proteins. Mol Cell 6, 11–22.

Teng, Y.S., Chan, P.T., and Li, H.M. (2012). Differential age-dependent import regulation by signal peptides. PLoS Biol 10, e1001416.

Toufighi, K., Brady, S.M., Austin, R., Ly, E., and Provart, N.J. (2005). The Botany Array Resource: e-Northerns, Expression Angling, and Promoter analyses. Plant J 43, 153–163.

Tsai, J.Y., Chu, C.C., Yeh, Y.H., Chen, L.J., Li, H.M., and Hsiao, C.D. (2013). Structural characterizations of the chloroplast translocon protein Tic110. Plant J 75, 847–857.

Vinti, G., Fourrier, N., Bowyer, J.R., and López-Juez, E. (2005). Arabidopsis cue mutants with defective plastids are impaired primarily in the photocontrol of expression of photosynthesis-associated nuclear genes. Plant Mol Biol 57, 343–357.

Vojta, A., Alavi, M., Becker, T., Hormann, F., Kuchler, M., Soll, J., Thomson, R., and Schleiff, E. (2004). The protein translocon of the plastid envelopes. J Biol Chem 279, 21401–21405.

Wu, F.H., Shen, S.C., Lee, L.Y., Lee, S.H., Chan, M.T., and Lin, C.S. (2009). Tape-Arabidopsis Sandwich - a simpler Arabidopsis protoplast isolation method. Plant Methods 5, 16.

Wu, G.Z., Meyer, E.H., Richter, A.S., Schuster, M., Ling, Q., Schottler, M.A., Walther, D., Zoschke, R., Grimm, B., Jarvis, R.P., and Bock, R. (2019). Control of retrograde signalling by protein import and cytosolic folding stress. Nat Plants 5, 525–538.

